# Quantifying pollen-vegetation relationships to reconstruct ancient forests using 19th-century forest composition and pollen data

**DOI:** 10.1101/039073

**Authors:** Andria Dawson, Christopher J. Paciorek, Jason S. McLachlan, Simon Goring, John W. Williams, Stephen T. Jackson

## Abstract

Mitigation of climate change and adaptation to its effects relies partly on how effectively land-atmosphere interactions can be quantified. Quantifying composition of past forest ecosystems can help understand processes governing forest dynamics in a changing world. Fossil pollen data provide information about past forest composition, but rigorous interpretation requires development of pollen-vegetation models (PVMs) that account for interspecific differences in pollen production and dispersal. Widespread and intensified land-use over the 19th and 20th centuries may have altered pollen-vegetation relationships. Here we use STEPPS, a Bayesian hierarchical spatial PVM, to estimate key process parameters and associated uncertainties in the pollen-vegetation relationship. We apply alternate dispersal kernels, and calibrate STEPPS using a newly developed Euro-American settlement-era calibration data set constructed from Public Land Survey data and fossil pollen samples matched to the settlement-era using expert elicitation. Models based on the inverse power-law dispersal kernel outperformed those based on the Gaussian dispersal kernel, indicating that pollen dispersal kernels are fat tailed. Pine and birch have the highest pollen productivities. Pollen productivity and dispersal estimates are generally consistent with previous understanding from modern data sets, although source area estimates are larger. Tests of model predictions demonstrate the ability of STEPPS to predict regional compositional patterns.

---

> It has been claimed that skilful hunters of old could bring their prey down even if they caught only a fleeting glimpse of its shadow. But still more remarkable appear the performances accomplished to-day by the pollen analyst. Out of pollen from crumbled clay or minute pieces of peat, taken from bits of earthenware or a stone axe, may be constructed a picture of the primeval forests which flourished in the region at the time when the pot or axe dropped, into the bog. This is, however, not the whole story.
>
> — Gunnar Erdtman, 1943

## 1 Introduction

Understanding changes in past forest composition provides valuable information about ecosystem response to biotic and abiotic factors at timescales outside the realm of most direct ecological observations. Fossil pollen records extracted from sediments of lakes and mires are a primary source of data about prehistoric forest composition (Brewer et al., 2012). Rigorous estimates of past forest composition from these records rely on ability to quantify the relationship between the composition of fossil pollen assemblages and the composition of the vegetation that produced them.

The pollen-vegetation relationship is governed by the processes of pollen production, transport, and deposition and has been the subject of empirical and theoretical study for decades (Davis, 1963; Tauber, 1965; Jacobson and Bradshaw, 1981; Jackson, 1994; Jackson and Lyford, 1999; Sugita, 2007*a,b*; Prentice, 1988). Pollen found in lakes and mires usually arrives by wind, coming from plants that vary in pollen productivity and dispersal properties. Some plant taxa are overrepresented in sedimentary archives while others are underrepresented or absent. Pollen-vegetation models (PVMs) are designed to make inferences about past vegetation composition or structure from fossil pollen data. Some PVMs employ multivariate methods to match pollen assemblages with vegetation types or biomes; these include modern-analog based approaches (Overpeck et al., 1985; Williams, 2003) and biomization (Prentice et al., 1996; Williams et al., 1998). Other PVMs relate abundances of individual plant taxa in vegetation with their abundance in pollen assemblages. These include pollen/vegetation ratios (R-values) (Curtis, 1959; Davis, 1963), linear-regression (Webb et al., 1981; Bradshaw and Webb, 1985), extended R-values (Parsons and Prentice, 1981; Sugita, 1994, 2007*a,b*), and, most recently, Bayesian hierarchical models (Paciorek and McLachlan, 2009; Garreta et al., 2010). In these models, pollen dispersal may be assumed uniform (Davis, 1963; Parsons and Prentice, 1981), modeled to decline away from the source as a simple inverse-square or other function (Webb et al., 1981; Calcote, 1995; Jackson and Kearsley, 1998), or parameterized using various pollen-transport models, usually based on the Sutton models of airborne particulate dispersal (Prentice, 1985, 1988; Jackson and Lyford, 1999; Sugita, 2007*a,b*).

PVMs are calibrated using pollen and forest data. Some PVMs, notably the LOVE/REVEALS family of models (Sugita, 2007*a,b*), require information about pollen dispersal properties for ndividual taxa, which are then used to parameterize diffusion-based models of particulate transport [e.g., Prentice (1985)]. In some cases, PVMs are calibrated against spatial networks of pollen assemblages extracted from surface-sediment samples that are cross-referenced to contemporary spatial datasets of vegetation (e.g., Webb et al. (1981); Williams (2003)). These cross-referenced pollen and vegetation datasets are then used to build empirical models that can be used to infer past forest composition from fossil pollen assemblages.

However, most models lack the ability to account for uncertainty in a formal, coherent way. Without some understanding of the uncertainty, it is difficult to make conclusions about changes across space and/or time. Our primary goal in this paper is to develop a PVM to estimate past vegetation composition and its uncertainty. As opposed to working with individual sites, we treat the data as a cohesive unit and allow it to directly inform us about changes in space and time. As in Paciorek and McLachlan (2009), we use a Bayesian hierarchical spatial approach (Spatio-Temporal Empirical Prediction from Pollen in Sediments, henceforth referred to as STEPPS) to model pollen counts from a network of ponds as a function of gridded forest-composition data, taking into account local pollen dispersal (i.e., pollen sourced from trees within a grid cell containing a pond), regional pollen dispersal (pollen sourced from trees in other grid cells), differential pollen production across tree taxa, and process uncertainty. The statistical model infers production and dispersal capability by finding parameter values that best explain the sediment pollen data at all sites given known vegetation across the domain, within the structure of how the model represents production and dispersal. Parameter estimates are used to estimate spatial maps of vegetation, and their uncertainty, even at locations without extant pollen data.

Most PVMs are designed to reconstruct vegetation for the relevant pollen source area of a depositional basin. Previous studies have shown that the size of the deposition basin may affect the relevant pollen source area, with sediment pollen from larger lakes providing better representation of the regional pollen signal (Jacobson and Bradshaw, 1981; Prentice, 1985; Sugita, 1994, 2007*a*). In the LOVE/REVEALS models (Sugita, 2007*a,b*), background pollen is estimated based on pollen counts from large lakes and is then used to inform local vegetation estimates from pollen counts at smaller depositional sites. STEPPS also divides pollen dispersal into local and non-local components. However, the operational definition of *local* varies among models, and within the literature. In Jacobson and Bradshaw (1981), local is defined as being within 10^2^ m of the depositional basin; in our case, local refers to the grid cell containing this basin, which is 10^3^ m - an order of magnitude larger.

Unlike many other PVMs, STEPPS assumes that depositional basin size does not affect the distance weighting of the surrounding vegetation. This is in part a consequence of the spatial resolution of the 8 km gridded vegetation data sets used here; our ability to resolve fine-scale compositional patterns near depositional basins is restricted by the coarseness of the available vegetation data. However, all deposition sites regardless of size receive much of their pollen from regional sources and hence are sensitive to broader-scale vegetation patterns and gradients (Bradshaw and Webb, 1985; Prentice et al., 1987; Jackson, 1991; Sugita, 2007*a,b*). Although the lack of higher-resolution vegetation-sampling likely imposes some error (Bradshaw and Webb, 1985; Prentice et al., 1987; Jackson, 1990), our goal is to predict vegetation at a regional scale and as such we do not expect to resolve fine-scale local heterogeneities.

Most spatially calibrated PVMs are not explicitly process-based, although sometimes the parameters in these empirical models give insight into process. For example, the y-intercept in linear regression models provides an estimate of pollen dispersed from outside the vegetation-sampling area (Webb et al., 1981). In contrast, Bayesian hierarchical models can be spatially calibrated and designed to include parameters that represent processes in pollen production, transport, and deposition (Paciorek and McLachlan, 2009). While both STEPPS and LOVE/REVEALS account for production and dispersal, neither model is fully mechanistic (e.g., explicitly based on particle-dispersal dynamics). The strength of the Bayesian PVM presented here relative to LOVE/REVEALS and other PVMs is its ability to estimate vegetation, process parameters, and uncertainty, by accounting for spatial dependence and borrowing information across space. We avoid the need for *a priori* parameter estimates (e.g., of pollen productivity, atmospheric conditions, or settling velocity) using a coherent hierarchical framework. Requiring parameter estimates is not an impediment per se, but some of the inputs required for other PVMs may be difficult to measure or quantify or variable through time and/or space (Jackson, 1994; Jackson and Lyford, 1999).

Extending PVMs to make inferences about palynological processes and vegetation composition for times other than the time of calibration requires the assumption that pollen-vegetation relationships are constant. This assumption of constancy seems reasonable at first approximation; although tree taxa may vary in their productivity and dispersal both in space and time, the relative differences among taxa should be less variable (Parsons and Prentice, 1981). In eastern North America and elsewhere, recent human land use has significantly altered forest composition and structure, indicating that this assumption of constancy in the pollen-vegetation relationship merits investigation (Kujawa et al., 2016). Most contemporary forests differ in density, biomass, size structure, and age structure from pre-settlement forests (Leahy and Pregitzer, 2003; Schulte et al., 2007; Rhemtulla et al., 2009; Goring et al., 2015b). Land clearance, forest management practices, and successional processes have led to fundamental changes in forest composition, and hence early-successional trees such as poplar and some maple species are more abundant now than in the 19th century (Thompson et al., 2013; Goring et al., 2015b). The overall pattern of forest mosaics in the region has also changed - at broad scales forests are more spatially homogeneous (e.g., forest management practices and recovery from land clearance) while at finer scales forests are more heterogeneous (e.g., locally idiosyncratic disturbance or land-use legacies) (Thompson et al., 2013; Wang, 2007; Li and Waller, 2015; Goring et al., 2015b). Finally, fire suppression during the 20th century may have led to increasing forest density and changes in abundance of fire-adapted species (Nowacki and Abrams, 2008). Critically however, the nature of change across the region varies; for example, fire suppression has led to higher biomass in northern Minnesota, while clearance and regeneration has resulted in lower biomass in southern Wisconsin and Michigan (Goring et al., 2015b).

These changes in forest composition, structure, and landscape pattern may have directly influenced pollen production and dispersal (Kujawa et al., 2016). Pollen production, for example, may be greater for trees growing in the open or at forest edges (Feldman et al., 1999). Pollen dispersal is affected by properties of both the forest canopy and the land surface (Jackson and Lyford, 1999). A more serious issue for pollen-vegetation relationships may be that at the landscape level, many contemporary forests have no historical analogues (Goring et al., 2015b). Shifting forest composition can alter pollen-vegetation relationships because of the Fagerlind effect, in which relative pollen representation of one taxon is affected by that of all other taxa contributing to an assemblage (Prentice, 1988). It is possible to account and correct for the Fagerlind effect using ERV models (Prentice and Webb, 1986; Jackson et al., 1995), but empirical corrections are contingent on particular vegetational contexts (Jackson and Kearsley, 1998).

Most PVMs are calibrated using contemporary forest data; few studies have examined settlement-era pollen-vegetation relationships (Schwartz, 1989). Although some have used historical forest cover maps to validate models based on modern relationships (Nielsen, 2004), a recent study has shown that pollen-based paleoclimate reconstructions calibrated with settlement-era pollen data differ significantly from those using modern pollen data (St Jacques et al., 2014) suggesting that shifts in pollen-climate relationships have occurred across the region. The compilation of settlement-era Public Land Survey (PLS) forest composition records from the western Great Lakes region (Bourdo, 1956; Schulte and Mladenoff, 2001; Almendinger, 1996; Liu et al., 2011; Goring et al., 2015b), along with a new dataset of settlement-era fossil pollen records from the upper Midwest (Kujawa et al., 2016), largely drawn from the Neotoma Paleoecological Database (www.neotomadb.org), provides an opportunity to calibrate a PVM for the early to mid-19th century, thus avoiding the effects of settlement, agriculture, and industrialization.

Cross-referencing settlement-era forest composition with appropriate fossil pollen data to construct a settlement-era pollen-calibration dataset requires identification of pollen samples that are contemporaneous with the settlement era. In eastern North America, land clearances during Euro-American settlement led to increases in certain agricultural and weedy plant taxa; these increases can be used as direct biostratigraphic markers of settlement in pollen records. Identifying the settlement horizon requires interpretation of pollen sequences; the uncertainty associated with this interpretation has not been assessed to date. Here we use expert opinion from four paleoecologists to reduce bias and quantify uncertainty associated with the identification of biostratigraphic signals.

Towards our goal of predicting vegetation and uncertainty, we develop and calibrate a Bayesian hierarchical PVM based on the STEPPS model (Paciorek and McLachlan, 2009). We calibrate the PVM using the settlement-era PLS forest composition and pollen datasets, and use these calibration results to predict settlement-era vegetation composition for the upper Midwestern US. We assess our ability to predict vegetation composition by comparing these spatial predictions with the PLS data. With both prediction and understanding in mind, we test variations of the calibration model against the data and compare results to our current scientific understanding of pollen processes. Specifically, we test and compare two standard isotropic pollen-dispersal kernels. We apply the PVM results to develop new estimates of relative pollen productivity and source area, and assess the relative importance of local and non-local pollen sources.

## 2 Data and Methods

### 2.1 Data

#### 2.1.1 Spatial domain

The study area is the upper midwestern US (UMW), including Minnesota, Wisconsin, and Michigan (see Fig. 1). This region includes two major ecotones: the “Tension Zone” between northern mixed forests and temperate broadleaved deciduous forests Curtis (1959), and the prairie/forest/savanna ecotone. Both of these ecotones have been heavily modified by contemporary land use (Goring et al., 2015b). We focus on the UMW because of the strong vegetation gradients associated with these ecotones and because of the dual availability of an integrated network of pre-settlement forest-composition estimates (Goring et al., 2015b) and high density of pollen sequences spanning the settlement era.

**Figure 1:**
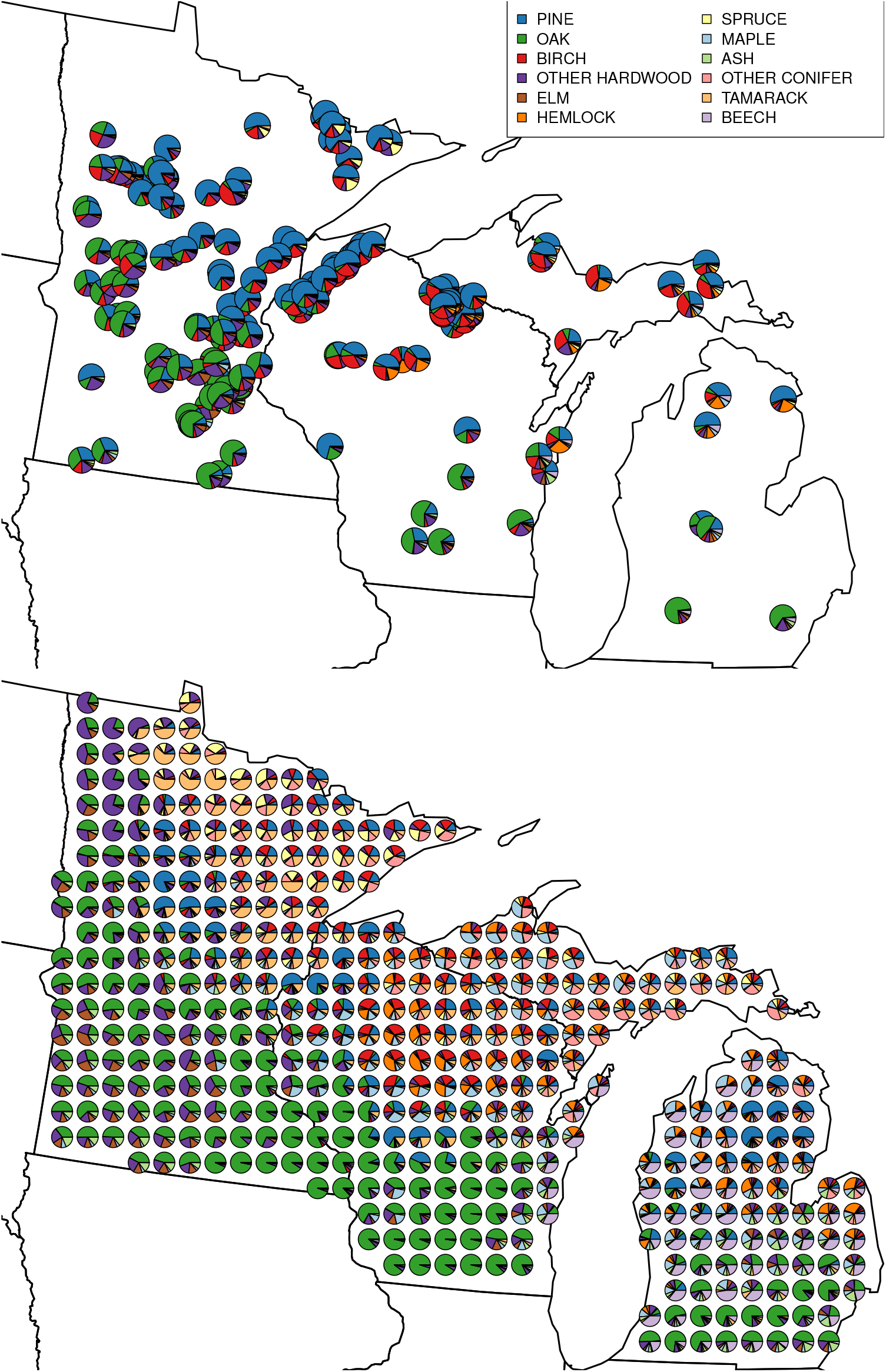
Pie maps depicting the relative composition of tree genera of pollen (top) and PLS vegetation (bottom) from the data. Note that the PLS data has been aggregated from 8 km to 32 km resolution for illustrative purposes.

#### 2.1.2 Public Land Survey (PLS) data

Prior to major European settlement, the US General Land Office conducted a Public Land Survey (PLS) throughout much of the United States to simplify the sale of federal lands. The PLS mapped the United States west and south of Ohio on a 1x1 mile grid for the sale of public lands (Stewart, 1935; White, 1983). Surveyors marked section corners with stakes, and recorded the closest two to four trees, measuring the distance from tree to point center, azimuth, tree diameter and the (often idiosyncratic) common name of these trees (Mladenoff et al., 2002).

Data from the PLS constitutes a systematic survey of the forest before settlement, and has been used by foresters, ecologists, and historians to learn about ecosystem and land-use change. In the UMW, the survey was conducted between 1834 and 1907 (Stewart, 1935).

Recently these point-level data have been aggregated across the UMW to a 64 km^2^ grid (8×8 km) on an equal-area Albers projection (proj4: +*init* =*epsg*:3175) and standardized to account for changing survey instructions through time (cf. Stewart, 1935) and to provide sufficient spatial resolution for regional-scale analysis (Goring et al., 2015b). This aggregation results in an average count of 134 trees per grid cell. This tree count per cell is low, but ecological patterns and regional structure of the dominant forest taxa are evident because of the uniforma spatial sampling (Goring et al., 2015b). In a given grid cell, the raw data provide noisy estimates of composition proportions because of the limited number of trees per cell. Instead, we use composition estimates provided by Paciorek et al. (2015), who applied a statistical model that borrows information across nearby grid cells to estimate the composition proportions in each grid cell with uncertainty. Here we work with these PLS composition estimates. These estimates are not data in the true sense, but assumed to be fixed for our purposes and are hereafter referred to as PLS data.

#### 2.1.3 Tree taxa

For the UMW domain, Goring et al. (2015b) report fifteen tree taxa, although the full dataset contains 28 taxa. Most taxa are resolved to the level of genera, and the list of taxa includes “No Tree” (21% of points) when surveyors did not find a tree within 300 links or 198 feet, “Unknown Tree” (0.001% of points) when the common name used to identify the tree could not be resolved, and “Other Hardwood” (0.02% of points) when the surveyed tree was clearly a hardwood, but could not be resolved beyond that level. We focus on a subset representing two functional groups (Deciduous and Coniferous taxa) composed of 10 tree genera that include the most abundant taxa as well as less-abundant but ecologically significant taxa. These taxa are: ash, beech, birch, elm, hemlock, maple, oak, pine, spruce, and tamarack (see Table 2 for scientific name translations). The remaining tree taxa, including “Unknown Tree”, are aggregated to “Other Hardwood” or “Other Conifer”, which amount to 10% and 7% of the included trees respectively. “No Tree” is excluded from further analysis.

#### 2.1.4 Pollen data

Advances in paleoecoinformatics (Brewer et al., 2012; Grimm et al., 2013) have made it possible to easily access and query large datasets using tools such as the multi-proxy Pliocene-Quaternary database Neotoma `(neotomadb.org).` The development of the Neotoma API `(api.neotomadb.org)` has provided a foundation for the development of a map-based search interface - the Neotoma Explorer (apps.neotomadb.org/explorer) - and the neotoma package for R (Goring et al., 2015a). Using these tools we identified 176 sedimentary pollen records in the Neotoma database from the UMW. To be considered for settlement-era calibration, records must contain a sample deemed representative of settlement-era vegetation; for this reason, we considered only records that included multiple pollen samples from the last 2000 years (under the assumption that one of these samples can be deemed representative). An additional 57 records, not available from Neotoma at the time of this project, were contributed by independent researchers (Kujawa et al., 2016). These additional records include 9 pollen-count time-series records and 48 records that consist only of the core-top and pre-settlement assemblages, hereafter referred to as before-after records. The dating controls available for the identified pollen records varied. To form our calibration data set we primarily relied on biostratigraphic markers of EuroAmerican land clearance. As a result radiometric and other dating controls were not explicitly considered during the pollen record screening process (but will be considered in the formation of the prediction data set). The new settlement-era pollen dataset is described in Kujawa et al. (2016).

Pollen records are obtained from morphological assignment of individual pollen grains from sediment samples into groups that correspond with plant taxa, which is a labor-intensive and skill-dependent process. Each pollen count represents a sample from a larger population (Maher et al., 2012; Maher, 1981), although uncertainty from pollen identification and counting is negligible for regional-to-global-scale syntheses focused on the most common taxa from lake and mire sediments (Webb et al., 1978*a*,*b*). These data are collected by independent researchers and as such require the standardization of pollen taxonomies to account for differing nomenclature and taxonomic resolution. Here pollen taxonomies were standardized using the STEPPS translation table from `neotoma`, which is reported as supplemental material.

#### 2.1.5 Elicitation of evidence for settlement in pollen records

Widespread land clearance during Euro-America settlement provided habitat for many ruderal plants, resulting in increases in non-arboreal taxa that are evident in pollen records (McAndrews, 1988). In the UMW, significant increases in ragweed *(Ambrosia*), docks and sorrels *(Rumex*), and grasses *(Poaceae*) typically mark this settlement horizon.

Here we use expert elicitation to identify the representative settlement horizon sample, defined to be the sample preceding the increase in agricultural indicator species, at each of the 185 sites with pollen count time series data. All expert elicitation relies on judgment, and there is often significant variability in expert’s determination of true signals (Bond et al., 2007, 2012). The degree of uncertainty resulting from expert identification of “settlement” likely varies among sites as a result of the temporal density of pollen samples, time-averaging of sediments, the rapidity of forest clearance and landscape transformation, the pollen representation of dominant trees, which can dampen or amplify the ragweed signal, and expert knowledge of the region and the late-Holocene history of the site. At some sites, the settlement horizon is unambiguous, while at others the transition is less clear (see App. A, Fig. A).

In the interest of reducing bias and quantifying uncertainty, we: 1) use consistent methodology to identify the settlement horizon in the pollen records, and 2) assess the variability in settlement horizon assignment among analysts. Four expert palynologists (Goring, Jackson, St. Jacques, Williams) participated in the elicitation exercise. Each expert was provided with pollen diagrams depicting proportional changes through time as a function of depth for the ten most abundant arboreal plus key indicator taxa for the last 2000 years as determined by the default Neotoma age-depth models. Experts were prohibited from relying on stratigraphic dates (radiocarbon or other) or age-depth model estimates of sample age. Experts were instructed to report the horizon as non-identifiable when they could not distinguish the settlement horizon, and to note the uncertainty of their settlement horizon assignments, with or without justification. Expert identifications were subsequently checked against age assignments based on age-depth models either obtained from Neotoma or constructed using original age controls and the Bacon age model (Blaauw and Christen, 2011). Results from this exercise were used to define depths associated with settlement horizon samples and their uncertainty. Pollen samples corresponding to the selected settlement horizon depths were included in our calibration data set. Expert elicitation results are included as supplementary material.

### 2.2 Models

In this section we describe the PVM. In our workflow, vegetation reconstruction is composed of two phases: calibration and prediction. In section 2.2.1, we describe the calibration model used to estimate the parameters that describe the processes that link vegetation composition to sediment pollen through pollen production and dispersal. Then, in section 2.2.2, we describe the prediction model used to estimate settlement-era vegetation composition. In section 2.3 we describe computational software and tools used to fit the models.

#### 2.2.1 Calibration model

Here we describe the calibration model used in the first phase of vegetation reconstruction. We first provide an overview of the model and then introduce the mathematical notation.

The observed pollen counts are a function of the vegetation on the landscape; i.e., the composition and abundance of trees on the landscape determines (in part) the composition and abundance of pollen deposited in the sediment. How much pollen the surrounding vegetation contributes is a complex function of forest characteristics (e.g., composition, structure, patchiness, spatial stationarity), climate (including wind), pollen productivity, grain size and morphology, topography, and proximity of the vegetation to a deposition basin.

We work on an 8 km grid defined by the resolution of the gridded PLS data and define each deposition site as having both local and non-local pollen contributions. Local contributions are made by the vegetation within the same grid cell as a site, and non-local contributions are made by vegetation from other grid cells. The relative contribution of local versus nonlocal pollen is determined by a model parameter, with the restriction that the sum of all contributions from the pollen source area must be one. We emphasize that here “local” refers to pollen from an area much larger than what would usually be referred to as such.

Contributions from non-local grid cells are not all equal; their relative contributions are distance-weighted according to a dispersal kernel, which determines how likely it is for pollen to travel from one grid cell to another. Dispersal is a well-studied phenomenon, and there is no consensus on a single best dispersal kernel; dispersal kernels vary in function complexity and flexibility, and some are better suited in certain cases than others (Clobert et al., 2012). Given this, we test two dispersal kernels, both of which have been used to model pollen dispersal (Clobert et al., 2012). The considered dispersal kernels are isotropic, meaning they depend only on the distance of the tree to the deposition site and not on the absolute position or direction. In reality, dispersal may not be isotropic, owing to prevailing wind directions during pollen-dispersal seasons, regional topography, surface roughness, and vegetation patchiness on the landscape. However, isotropic dispersal kernels are a reasonable and widely used approximation (Sugita, 2007*a,b*), because turbulent atmospheric mixing below the boundary layer tends to smooth out anisotropies. We define a maximum dispersal distance that determines an upper limit to the distance that pollen is able to disperse from the source; this reduces computation without a significant loss of information.

Here, we choose two kernels that are representative of the short-and long-tail kernel classes, the Gaussian kernel (GK) and the inverse power-law kernel (PLK), and compare between these generalized functional forms. The GK is often used as a reference against more leptokurtic kernels such as the PLK and can represent dispersal through diffusion. While the GK has a simpler mathematical representation, fatter-tailed kernels (such as the PLK) in general perform better than those with thinner tails (like the GK or Exponential) when modelling pollen dispersal (Devaux et al., 2007; Austerlitz et al., 2004). However, pollen dispersal data needed to validate pollen dispersal models are generally unavailable, especially at the large spatial scales required to adequately test long-tailed kernels (Clobert et al., 2012).

To determine how much pollen a grid cell contributes to a deposition site such as a pond, the vegetation composition proportion vector is first scaled by taxon-specific parameters that account for differential pollen production among tree species. The local contribution is determined by the scaled vegetation vector for the grid cell that contains the pond. For all grids cells in the domain that do not contain the pond, their scaled vegetation vectors are multiplied by the respective weights assigned by the dispersal kernel (which depend on the distance to the pond and the kernel parameters). The total non-local contribution is the sum of the scaled and weighted vegetation vectors over all grid cells that do not contain the pond. The relative importance of the local and non-local contributions is determined by the local strength parameter, and the sum of the local and non-local contributions determines the predicted sediment pollen at the pond.

##### Calibration model details

Here we give a mathematical description of the calibration model. As described, the spatial domain is regular grid composed of 64 km^2^ discrete cells, but the underlying vegetation composition and dispersal spatial processes are continuous functions of space. Spatial cells are indexed by *s* = 1, …, *S*, and for the full UMW spatial domain *S* = 8013. All grid cells in the domain are potential sources of pollen.

Pollen produced by vegetation within each grid cell can be deposited within that grid cell or dispersed into the neighborhood around that grid cell. For grid cell *s_i_*, the pollen produced by taxon *p* within that cell that remains local is given by

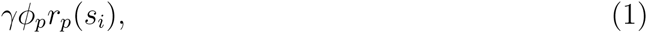

where *γ* is the proportion of pollen produced in *s_i_* that is deposited in *s_i_, ϕ_p_* is the scaling factor that accounts for differential production, and *r_p_(s_i_)* is the proportional abundance of vegetation of taxon *p* in *s_i_*.

The remaining 1 − *γ* proportion of pollen produced in *s_i_* is dispersed to other grid cells according to an isotropic dispersal kernel centered at *s_i_*. The dispersal kernel weights all pollen dispersing away from the focal cell as a function of the distance from *s_i_* to any neighboring cell *s_k_* by *w(s_i_, s_k_)*.

The two dispersal kernels considered are the GK, defined as

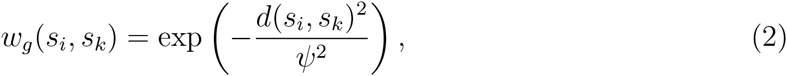

where *d(s_i_,s_k_*) defines the distance between cells *s_i_* and *s_k_* and *ψ* > 0 is a parameter that describes the spread of the kernel, and the PLK, defined as

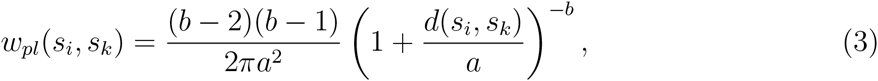

where *a* > 0 and *b* > 2 are parameters that determine the kernel shape. As expected, the weight assigned by both kernels is a decreasing function of distance - less pollen is distributed farther away.

Pollen produced by taxon *p* dispersing from *s_i_* to *s_k_* is then calculated as

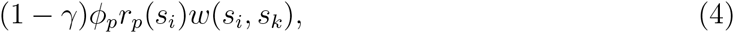

where *w* ∈ {*w_g_, w_pl_*}.

So far we have characterized the model from a source-based perspective, describing how pollen disperses from the grid cell in which it was produced. To quantify vegetation we need to interpret the counts of pollen grains from the sedimentary record from site *i* in grid cell *s_i_*. These pollen grains were produced in and dispersed from grid cells within the maximum dispersal distance, and have been deposited, preserved, and counted. The pollen arriving at site *i* is equal to the sum of the locally deposited pollen plus the pollen dispersed to *s_i_* from other grid cells in the domain that are within the maximum dispersal distance of *s_i_*. Therefore the pollen of taxon *p* arriving at pond *i*, referred to as 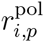, is given by

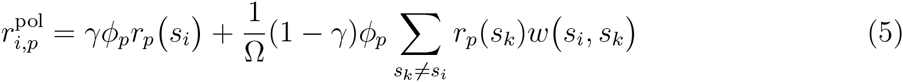

where Ω is a normalizing constant equal to the sum of the weights of all the cells to which pollen can be dispersed, defined to be a circle of radius 700 km. Note that Ω depends on the dispersal kernel, and therefore varies as a function of dispersal kernel parameters. However, for a given a set of parameter estimates, Ω is constant across the domain.

Finally, we need to relate the arriving pollen 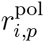 quantified in Equation 5 to the pollen count data. To do this, we consider that the pollen data are overdispersed - the counts are more variable than we might expect if they were multinomially distributed. This overdispersion is at least in part (and perhaps mostly) attributed to heterogeneity in the pollen production and dispersal processes that is not accounted for in our mathematical characterization but is also an artifact of the comparison of pollen counts at a point location (the depositional site) situated anywhere within a focal grid cell to a grid-based estimate of pollen deposition. Therefore, we model the pollen counts at depositional site *i*, denoted by ***y**_i_* = (*y*_*i*1_,…, *y_iP_*), with a Dirichlet-multinomial (DM) distribution, a compound distribution used to account for overdispersion in multinomial count data (that simplifies to the multinomial distribution in the case of no overdispersion). We have that

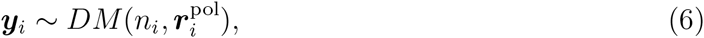

where 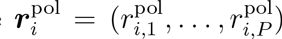 and *n_i_* is the number of pollen grains counted at site *i*. The precision *α_i_* of the distribution is equal to the sum of Equation 5 over all taxa

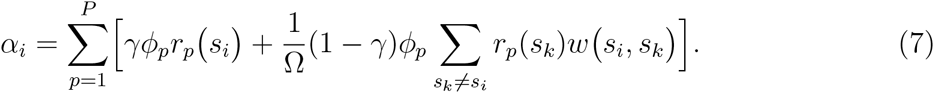

Note that the precision is affected by the proximity of a depositional site to the domain boundary. We do not have PLS data outside of the defined UMW domain, so we cannot quantify the pollen dispersed from vegetation outside the domain arriving at depositional sites in our domain. The repercussion of this is that the sum of weights of the grid cells contributing pollen to sites close to the domain boundary is less than the sum of weights of the total potential contributing neighborhood, defined as Ω. The result of having a smaller contributing neighborhood is that the local pollen contribution is effectively up-weighted relative to the non-local contribution.This edge effect will have the greatest impact at sites that are close to the boundary and near regions of vegetation that contribute pollen which are not accounted for (some sites are near the boundary but are surrounded by lakes which do not contribute tree pollen). The northeastern tip of Minnesota is likely the most affected part of the domain; it likely receives pollen from Canada where we do not have vegetation data. However, as long as there are sufficiently many sites for which we do have vegetation for most of the source area then these edge effects will not significantly influence our process parameter estimates (process parameters are not spatially-dependent).

##### Model priors

Non-informative priors are assigned to model parameters. For both dispersal kernels, we set the priors *ϕ_p_* ∽ uniform(0, 300) for *p* = 1,…, *P* and *γ* ∽ uniform(0,1). For the GK we define the prior *ψ* ∽ uniform(0, 2), while for the PLK we have *a* ∽ uniform(0,500) and *b* ∽ uniform(2, 6). Note that we work in megameters (Mm), where 1 Mm = 1000 km.

##### Model variants

As currently described, neither dispersal kernel (GK or PLK) nor the parameter *γ* vary by taxon. In reality pollen dispersal varies among species; pollen-grain morphology, size, weight, and release mechanism all vary systematically among plant taxa (Jackson and Lyford, 1999). We let the data inform us if there is sufficient evidence to support taxon-specific dispersal parameters by testing alternative models with exchangeable priors (Gelman et al., 2014*a*). An exchangeable prior defines the same prior distribution on each of the *P* parameters in question as a function of shared hyperparameters. When the variance hyperparameter is estimated to be small, there is not sufficient evidence in the available data to support taxon-specific parameters.

In the GK case we use the exchangeable priors

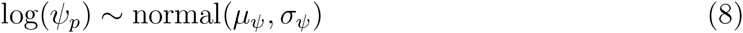

for taxon *p* = 1,… , *P* where *μ_ψ_* ∽ uniform(log(0.1), log(2)) (the log of the *ψ* prior interval in the single-taxon case), and *σ_ψ_* ∽ half-Cauchy(0, 2), a half-Cauchy with mean of 0 and scale of 2 (Gelman et al., 2006). For the PLK case, when *a* and/or *b* are allowed to vary their exchangeable priors are analogous to the exchangeable prior defined for *ψ* (see Equation 8). Note that for the PLK, *b* is allowed to vary by taxon only when *a* varies by taxon.

When *γ* is allowed to vary by taxon, we define a logit-normal prior such that the probability density function for 0 ≤ *γ_p_* ≤ 1 is given by

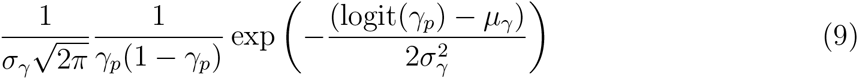

where *μ_γ_* ∽ uniform(—2, 2) and *σ_γ_* ∽ half-Cauchy(0, 5).

Allowing *γ* and/or the kernel to vary by taxon results in a total of 10 model variations, described in Table 1.

**Table 1:** Goodness-of-fit measures for the calibration and prediction models for model variants. For the calibration model, we report the Wantanabe-Akaike Information Criterion (WAIC). The exchangeable priors indicated that the data do not support a taxon specific *b* in the PLK model; for this reason we do not report the WAIC for the models with taxon-specific *b*. Overall, calibration models with the PLK have lower WAIC values. For the prediction model, we report the dissimilarity between the PLS data and the predictions as measured by the sum of the cell-wise Euclidean distances.

| Kernel | Model | WAIC | Dissimilarity |
| --- | --- | --- | --- |
| Gaussian | Base | 12966 | 2062 |
| Gaussian | Variable $\gamma$ | 12899 | - |
| Gaussian | Variable $\psi$ | 12895 | - |
| Gaussian | Variable $\psi$ and $\gamma$ | 12804 | 2078 |
| Power-law | Base | 12693 | **1979** |
| Power-law | Variable $\gamma$ | 12630 | - |
| Power-law | Variable $a$ | 12563 | - |
| Power-law | Variable $a$ and $b$ | - | - |
| Power-law | Variable $a$ and $\gamma$ | **12548** | 2044 |
| Power-law | Variable $a$, $b$ and $\gamma$ | - | - |

##### Model comparison

To reduce the number of candidate models, we first assessed the exchangeable prior variance estimates. When there was insufficient evidence to support the case for taxon-specific parameters, as indicated by (i) an estimated variance close to zero and (ii) overlapping credible intervals for the posterior distributions of the taxon-specific parameters, the model was no longer considered a candidate model.

Then, to determine the kernel and model that best fit the data we performed a formal model comparison using the Watanabe-Akaike Information Criterion (WAIC) (Watanabe, 2010). The WAIC provides a relative measure of loss of information (as does the more familiar AIC) in a Bayesian context, and accounts for the full posterior distribution as opposed to conditioning on a single point estimate (as in AIC and DIC) (Gelman et al., 2014*b*). One of the benefits of the WAIC is that it avoids the necessity of knowing the number of model parameters (as required by AIC).

##### Assessing productivity and dispersal estimates

To assess our level of agreement with our current scientific understanding of pollen dispersal and production processes, we compare our productivity and dispersal estimates to previously published results. We do not expect agreement among absolute quantities; if STEPPS is well-defined, we expect the relative ordering of taxa with respect to both of these processes to be consistent (at least in general) with our current understanding.

In previous pollen-vegetation studies, pollen productivity was quantified in several ways. In models that assume a linear relationship between pollen and vegetation proportions (Webb et al., 1981; Bradshaw and Webb, 1985; Jackson, 1990), the slope coefficient accounted for differences in pollen productivity among taxa but also included effects of dispersal. The degree to which these effects are confounded in the slope coefficient depends on the nature of the vegetation sampling; in general, the larger the area of sampled vegetation, the more likely it is that productivity and dispersal are confounded. However, the size of the sedimentary basin also affects our ability to distinguish these effects. In the Extended R-value (ERV) model, deposited pollen is a function of nearby vegetation composition, plus a background pollen source (Prentice and Webb, 1986; Prentice et al., 1987; Marquer et al., 2014; Broström et al., 2008). The α-slope coefficient that relates deposited pollen from nearby sources to nearby vegetation measures productivity, but as in the linear regression model, it is also confounded by dispersal processes to some degree. With the goal of quantifying unconfounded pollen productivity, pollen dispersal biases can be removed from the α-slope coefficients estimated from the ERV model (Sugita et al., 1999). We compare our productivity estimates *ϕ_p_* to pollen productivity estimates (PPEs) determined by each of the above methods. For ease of comparison, each set of PPEs is rescaled to the [0,1] interval.

Comparing pollen dispersal estimates with our established understanding of dispersal is less straightforward. In STEPPS, dispersal is defined as the sum of locally- and non-locally-deposited pollen, where non-local pollen is distance-weighted by a dispersal kernel. As such, there is no single parameter that quantifies dispersal. However, we can compute a capture radius, i.e., the radius of the circle within which some proportion of the total released pollen is deposited. We use this measure as an crude approximation of distance travelled, which is inversely related to falling speed, at least for a fixed release height, wind-speed, and degree of atmospheric stability.

#### 2.2.2 Prediction model

Most PVMs are developed with the goal of making predictions of vegetation composition from pollen data for past times. The PVM here is our first step towards that goal; in future work it will be used to reconstruct vegetation for the UMW for the last 2000 years. For now, we assess the predictive ability of the PVM by predicting the settlement-era vegetation used in calibration. This allows us to evaluate the PVM and the effects of the model variants on composition.

The prediction and calibration models are similar, in that the relationship between vegetation composition and sediment pollen is the same. Now we would like to make predictions in the absence of composition data, using the calibration process parameter estimates. The vegetation composition now becomes a vector of unknown parameters, with some defined spatial structure. This defined spatial structure is determined by the vegetation data used in calibration and constrains the estimated composition by quantifying the smoothness of compositional changes across the landscape.

More specifically, the structure of the relationship between the pollen counts and vegetation is identical to Equations 5 & 6. The underlying composition, unknown from the perspective of prediction, and denoted by *g*, is modelled as a Gaussian process, where for the set of points in our domain, defined by the grid cell centers, *g* follows a multivariate normal (MVN) distribution where

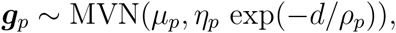

where *d* is the matrix of distances between grid cells, *η_p_* and *ρ_p_* are taxon-specific spatial covariance parameters, and *μ_p_* is the taxon-specific mean. The covariance paramaters *η_p_* and *ρ_p_* are estimated *a priori* by fitting a Gaussian process to the settlement-era composition data. The proportional vegetation *r_p_(s)* is linked to the underlying corresponding spatial process *g_p_(s)* through an additive log-ratio sum-to-one constraint,

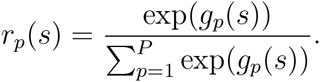

The full prediction model (not described here) used to estimate composition at multiple times accounts for covariance across both time and space.

Estimating the underlying vegetation process at each spatial location in the UMW domain is computationally challenging, at least for a grid of the specified resolution. To overcome the burden of long computational times, we use a Gaussian predictive process model in which the process is estimated at specified knot locations and then scaled up to the full domain (see Finley et al. (2009)). Here we use 120 knot locations, whose locations are determined using the kmeans clustering algorithm (MacQueen et al., 1967).

Settlement-era vegetation composition is estimated for the subdomain composed of Minnesota, Wisconsin, and the Upper Peninsula of Michigan using the settlement-era sediment pollen calibration data set. The Michigan Lower Peninsula is disjoint from the remainder of the domain, and only a few pollen samples are available for this region, and it is therefore excluded from the prediction domain in this analysis.

The dissimilarity between the predicted composition for different underlying calibration models and the PLS data is determined by measuring the Euclidean distance between the two sets of multi-category outcomes at each grid cell. More specifically, we compare the settlement-era PLS data ***r**(s_i_)* with the posterior mean of the composition estimates obtained from the prediction model 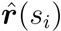. Then, for location *s_i_* the dissimilarity is

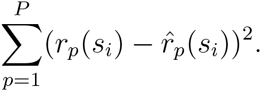

This measure is analogous to the Brier score, which measures the accuracy of probabilitic predictions (Gneiting and Raftery, 2007). Total dissimilarity is obtained for each set of predictions by summing the dissimilarity values over all locations.

### 2.3 Numerical implementation

We cannot directly sample from the joint posterior, so we rely on MCMC methods. With the goal of achieving more efficient sampling with respect to the effective sample size per unit time, we used the Stan v2.6.2 statistical modeling software to estimate parameters (Stan Development Team, 2014). Stan uses the No-U-Turn Sampler, a variant of the Hamiltonian Monte Carlo method which is a gradient-based sampling method that uses the directional derivatives from the gradient to make informed decisions about how to move along (and sample from) the joint posterior surface (Hoffman and Gelman, 2011).

The complex nature of the prediction model in combination with the domain size make it prohibitively slow to estimate prediction model parameters. To overcome this issue, we modified the Stan software to have it call a function that analytically computes the model gradient, thus avoiding our reliance on automatic differentiation. This modification allowed us to use parallel computation via openMP (OpenMP Architecture Review Board, 2008).

Ten candidate calibration models are considered: four GK and six PLK variants. For each model, we ran three chains using different random seeds and initial conditions with a warm-up of 250 iterations, followed by a sampling period of 1700 iterations. For the prediction models, we ran a single chain with a warm-up of 150 iterations followed by a sampling period of 1000 iterations. For all models, warm-up iterations were discarded and not used in further analysis. Convergence was assessed using a potential scale reduction factor and an effective sample size estimator (Gelman et al., 2014*a*). All scripts are made publicly available on github (https://github.com/PalEON-Project/stepps-calibration).

## 3 Results

### 3.1 Elicitation of evidence for settlement and site suitability

All 185 sites with pollen-count time-series were considered; only the before-after sites were omitted from this exercise. For 59 out of 185 sites the four experts were in complete agreements: they all identified the same settlement horizon (53 sites) or agreed that no such event could be identified (6 sites). The remaining sites varied in level of disagreement - for 79 sites, experts identified two settlement horizons; for 39 sites 3 were identified; and for 8 sites there was no agreement. These results confirm our hypothesis that identification of biostratigraphic horizon is subject to variability among analysts. To our knowledge this exercise is the first to quantify this uncertainty.

Of the six sites that experts agreed had no discernible settlement signal, two were from Rice Lake, which is an anomalous site locally dominated by wild rice *(Zizania*), with very high grass *(Poaceae*) pollen representation (McAndrews, 1969); it lacks any clear indication of settlement. The remaining four included: Rossburg peat bog in Minnesota (Wright et al., 1969); Disterhaft Farm Bog in Wisconsin (Webb and Bryson, 1972); Kirchner Marsh in Minnesota (Wright et al., 1963); and Stewart’s Dark Lake in Wisconsin (Peters and Webb, 1979). Three of these four sites are bogs and marshes with local marshland non-arboreal plants growing at or near the coring location; these marshland plants may overwhelm the signal from agricultural non-arboreal indicators.

Based on these results, we were faced with the challenge of establishing criteria to determine which sites to include in the calibration data set and where to place the settlement horizon for each site. Ideally, the now quantified settlement horizon uncertainty would be included into the modelling framework. In favor of simplicity, for now we use a single settlement horizon for each suitable site.

Sites were considered unsuitable if: 1) a majority of experts (more than half) chose not to assign a settlement horizon (12 sites), or 2) if half of the experts chose not to assign a settlement horizon and the other half identified settlement horizons whose estimated ages were far from the approximate time of settlement (7 sites). For this second criterion, settlement horizon sample ages from existing age-depth models were compared to 1850 AD, an approximate year of settlement in the UMW. Modeled ages were defined to be far from the expert-identified settlement horizon if discrepancies were greater than 500 years. All 19 unsuitable sites were excluded from further analysis.

Pollen records were checked for their representativeness and possible identification errors. To reconstruct regional vegetation, we would like pollen records to contain some regional signal. However, some sites were likely sampled as a result of their (relatively speaking) unusual composition or local setting, and may lack a strong regional signal. We identified two anomalous sites: Ocheda Lake and Tamarack Creek. Ocheda Lake is a large but shallow glacial kettle lake in Southwestern Minnesota (Neotoma site 6507). Pollen diagrams indicate a relative spike in red maple *(Acer rubrum*) pollen in the expert-identified settlement horizon sample. This may represent a misidentification given that red maple is rare in southwestern Minnesota. It is possible that this pollen comes from box elder maple (*A. negundo*) or silver maple (*A. saccharinum*), which occur along streams and lakes. We assume that this pollen is from the maple genus, and that the high count is a result of a maple anther falling into the sample. Hence, at this site we set the maple pollen count to zero. Tamarack Creek is a tamarack bog situated in an unglaciated wetland in the Driftless Area of western Wisconsin (Davis, 1979). This site is characterized by unusually high tamarack pollen counts for the top 80 cm of sediment. In particular, the settlement horizon sample is composed of 40% tamarack, while all other UMW settlement-era pollen records contain a maximum of 1% tamarack. Tamarack trees and pollen are known for being locally concentrated in bogs as a result of poor pollen production and dispersal. We set the tamarack count to zero to remove the local signal and use the remaining counts to represent composition from beyond the bog. We accept that these decisions may mask some true local-scale heterogeneity within the regional vegetation.

After completion of the elicitation exercise, a follow-up examination revealed several cores with top samples that were older (and deeper) than expected. Typically the age of a true core top corresponds with the year of sampling (with some margin of error). Further investigation uncovered that five of the cores included in our analysis were missing core tops. (The soupy sediments just below the sediment surface are often difficult to collect, and, depending on time, equipment, field conditions, and study purpose, investigators sometimes do not recover these upper sediments intact.) These errors are a result of data errors in Neotoma and the confusion associated with reporting depths as being either the depth below the lake surface or sediment surface. For Lake Mary (Webb, 1971), Green Lake (Lawrenz, 1975), Chippewa Bog (Bailey and Ahearn, 1981), and Ryerse Lake (Futyma, 1982), we determined that the core tops pre-date settlement, and these sites were therefore discarded. A third site, Lake Kotiranta, had a core top previously identified as a representative pre-settlement sample, and this sample was retained for inclusion in the calibration data set (Wright et al., 1969). The absence of core tops at these sites have been reported to Neotoma and the records amended accordingly. For each of these cores, 1-3 experts identified a pre-settlement sample (although to be fair, they made this judgment based solely on the stratigraphic diagram and no information about site characteristics, and likely assumed that the uppermost sample was in fact a surface sample).

After the suitability screening, we were left with 164 time series of pollen counts, and the question of how to assign the best estimate of pre-settlement. Given the sample resolution and an even number of experts, neither the mean nor the median were options. Keeping in mind that we would rather err on the side of caution and choose a depth that was more certainly pre-settlement (deeper) than one that represented post-disturbance times (more shallow), we ordered the expert-identified sample depths from deepest to shallowest, and selected the second in this list (typically the second deepest, although sometimes the deepest when there was some agreement among experts). The 48 records that consist of only a core top and previously identified settlement horizon sample were not candidates for the elicitation exercise. The decision to include these additional data in the calibration data set was debated - the settlement horizon samples for these cores were identified using different methodology (and analysts) than the remaining sites. In the end, we opted to include these samples based on our confidence in the data set and analysts and the recognition that including these sites would result in a substantive increase in sample size. In total we had 213 settlement-era sites in the UMW calibration data set and 21 sites considered inadmissible (Table C).

### 3.2 Exploratory data analysis

To visually assess the relationship between sediment pollen and tree taxa, we compare pie maps that depict the relative proportions of taxa across space (Fig. 1). In these maps the broadscale patterns of pollen and PLS data are consistent, both capturing the major regional ecotones. However, there are striking differences. First, pine dominates the pollen records for most of the northern half of the domain. However, in the PLS data pine proportions are occasionally high but the average proportion of pine across the domain is low (< 20%).

Second, in the vegetation composition data, some regions that have higher relative abundances of hemlock, tamarack, and maple are not apparent or much less pronounced in the pollen data. These taxa, particularly tamarack and maple, are generally underrepresented in pollen assemblages (Webb et al., 1981; Bradshaw and Webb, 1985; Jackson, 1990). As expected, these maps confirm that pollen data can detect key vegetational gradients, but that the relationship between pollen and vegetation is complex and taxon-dependent.

To further assess spatial distributions of pollen versus PLS data, we plot the data as heat maps by taxon. Differences in extent of these distributions indicate pollen dispersal beyond species range limits. For example, for both birch and pine, which are overrepresented in pollen records (Webb et al., 1981; Bradshaw and Webb, 1985; Jackson, 1990; Williams and Jackson, 2003), the distributions of sediment pollen extend well beyond the range boundaries of these taxa shown in the PLS data (Fig. 2, columns 1 and 2).

**Figure 2:**
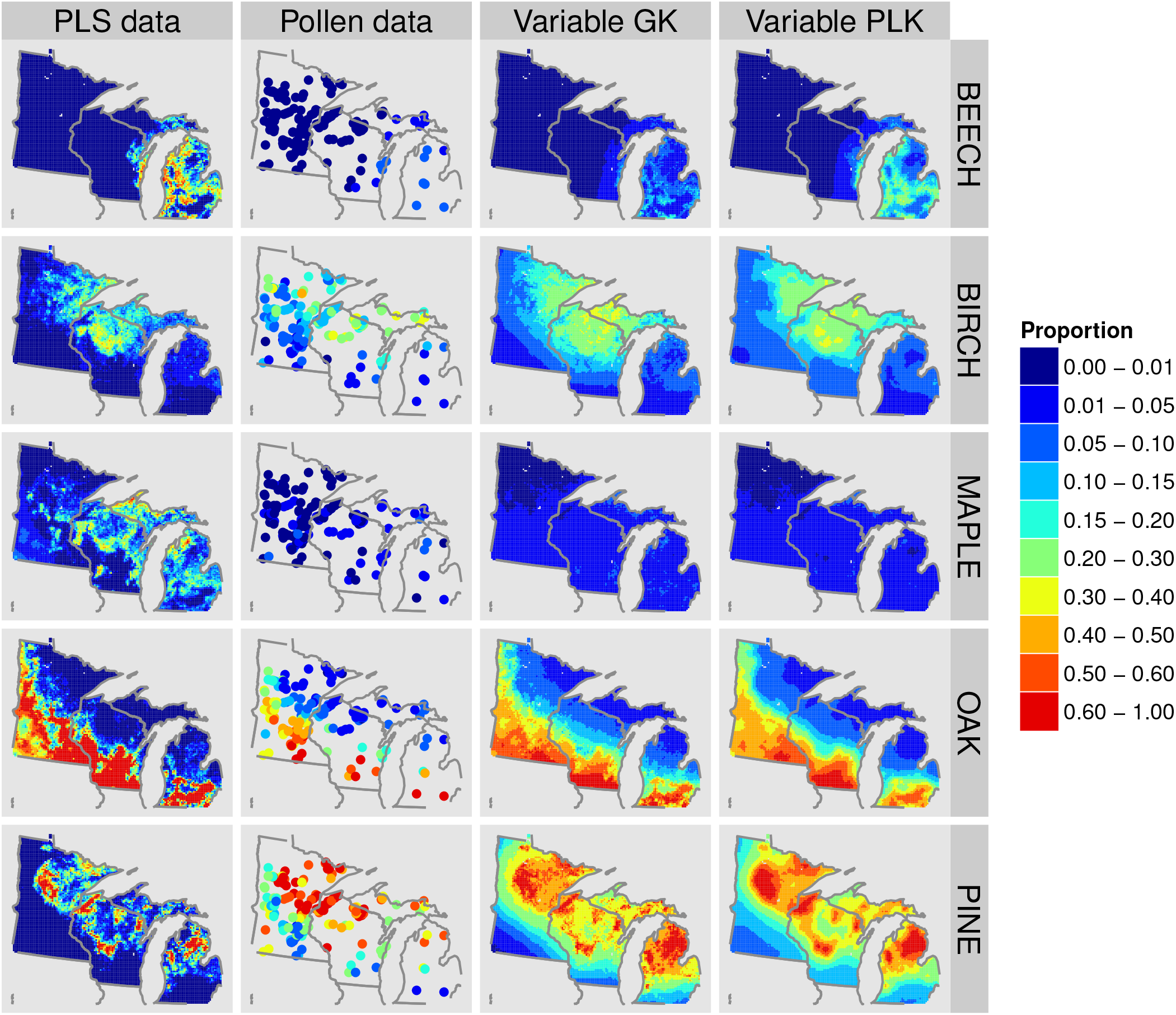
Heat maps of the PLS data, pollen data, and predicted sediment pollen for the variable Gaussian (GK) and power-law (PLK) kernel models for a subset of modelled taxa.

### 3.3 Modelling results

For the ten considered calibration models, effective sample sizes of the dispersal and production model parameters ranged from 1954 to 6000 for the GK models, and 480 to 6000 for the PLK models. In all ten cases, the three chains converged to the same posterior distributions. Trace plots and the potential scale reduction factor illustrate the efficient mixing achieved by the sampler (not shown).

To compare among models, we assess the exchangeable priors hyperparameter estimates. All parameter and hyperparameter estimates support taxon-specificity, except for the PLK models that allow *b* to vary by taxon. For these models, the estimated variance for the exchangeable prior for log(*b*) is small (9 · 10^−6^ for the variable *a*, and *b* case; 1 · 10^−4^ for the variable *a, b*, and *γ* case). In addition, the 95% credible intervals for the estimate of *b_k_* for all *k* overlap, supporting the conclusion that the values of *b_k_* are not statistically different. In light of this, we omit these two PLK variable *b* models.

For the remaining eight candidate models (four GK and four PLK variants) we compare the WAIC (Table 1). According to the WAIC the PLK outperforms the GK - even the most flexible GK model (variable *ψ* and *γ*) has a WAIC that is higher than that of the least flexible PLK model (base model). The PLK model with variable *a* and *γ* results in the lowest WAIC, indicating that it performs the best out of the suite of considered models.

In subsequent analyses, we compare the two base models (GK and PLK), and the models with the lowest WAIC for each kernel type - the GK variable *ψ* and *γ* model and the PLK variable *α* and *λ* model (hereafter referred to as the variable GK and PLK models).

In Fig. 3, the raw pollen proportions are plotted against the vegetation proportions for each grid cell (red crosses). The pollen-vegetation relationship is clearly not 1:1. In particular, some taxa, such as beech, maple, other conifer, and tamarack, are underrepresented-they can appear in large proportion on the landscape, but never in equally large proportion in the pollen record. Some relationships are more difficult to characterize. For example pine varies from sparse to abundant on the local landscape, but is consistently abundant in pollen assemblages in the northern half of our study area, regardless of local tree abundance.

**Figure 3:**
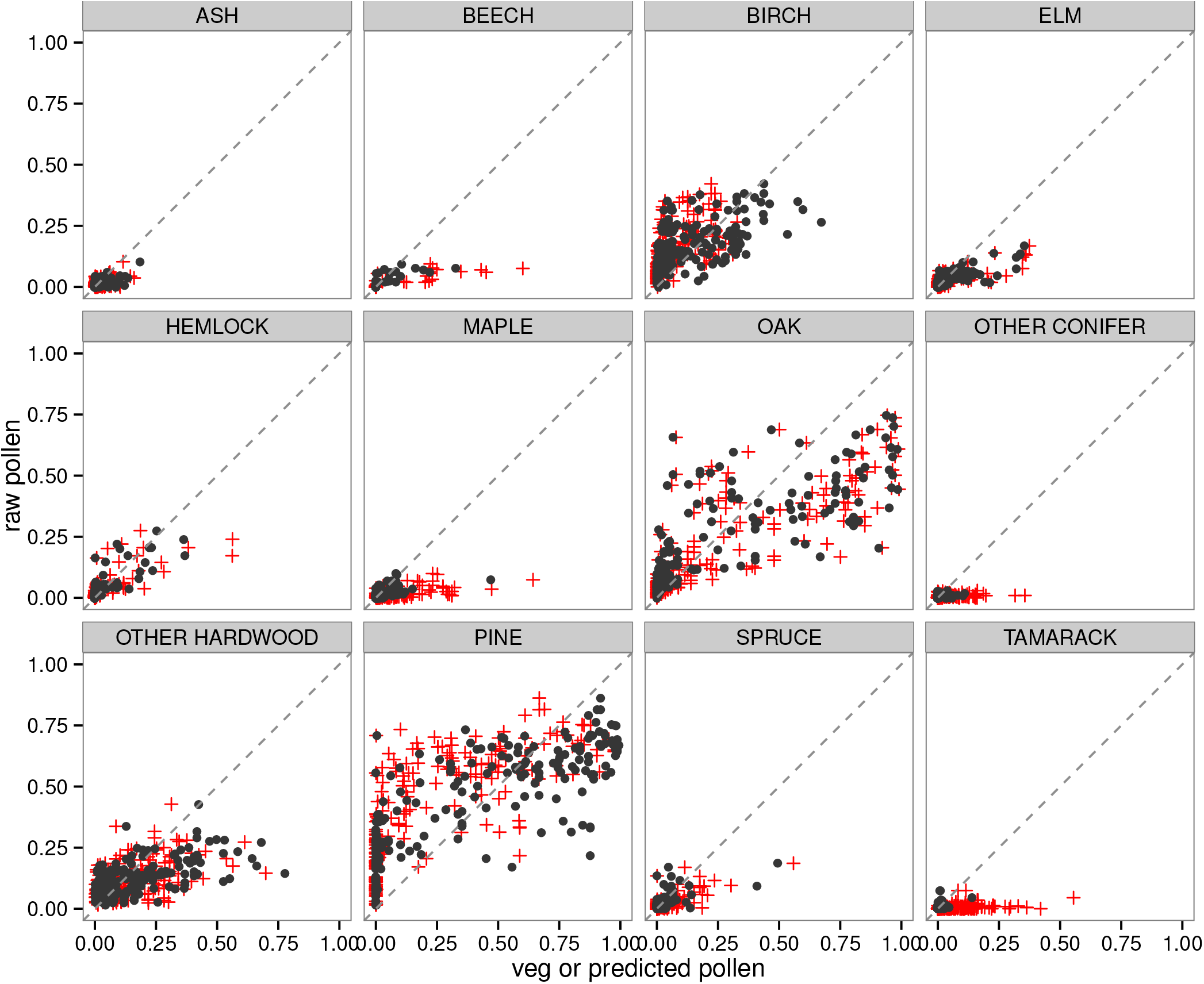
Pollen proportions plotted against local vegetation proportions (red crosses) or local vegetation proportion scaled by the pollen production factor *ϕ_k_* for the variable power-law kernel (PLK) model (black dots).

These relationships are complicated in part because of differential pollen production, which the calibration model accounts for with the taxon-specific scaling parameter *ϕ_k_*. After scaling the local vegetation in a cell by *ϕ_k_*, we compare the raw pollen proportions with the phi-scaled vegetation (Fig. 3; black dots). Here we show the scaled vegetation for the variable PLK model, but the results are similar for the variable GK and base models. Scaling by *ϕ* does improve our pollen estimates; for many taxa, the black dots have shifted towards the 1:1 line. This is especially true for maple, tamarack, and other conifers, suggesting that these pollen-vegetation relationships depend primarily on pollen production from local sources, and that dispersal for these taxa is limited. This finding is consistent with results of Jackson (1990, 1991) and others, which showed that pollen dispersal as well as productivity play important roles in their underrepresentation in pollen assemblages. However, for most of the remaining taxa the improvement is minimal, indicating that simply scaling the local vegetation proportions is not sufficient to predict deposited pollen.

In Fig. 4, we plot the raw pollen proportions against both the vegetation (red crosses) and predicted pollen for the variable PLK model (black dots). In this case the predicted pollen is based on the local plus non-local predictions obtained from the full calibration model. Now that non-local dispersal has been accounted for, the raw versus predicted pollen points fall more closely along the 1:1 line (compare the black dots in Fig. 3 to those in Fig. 4; improvement most noticeable for oak and pine). This indicates that, at least in our domain, non-local dispersal processes account for much of the pollen that arrives at any given location. For most taxa, the improvement that we gain by using more flexible models with taxon-specific parameters versus the base models is not qualitatively obvious (not shown). However, for hemlock, there is an obvious improvement in our ability to predict sediment pollen with the more flexible models. Both base models underpredict hemlock pollen at sites where there are relatively high proportions of hemlock (although hemlock pollen never exceeds 28% at any site in our domain), but this is not the case for the variable GK and PLK models. This suggests that with the base models the relative contribution of local versus non-local hemlock pollen is estimated to be less than the true value, i.e. *γ* is too small.

**Figure 4:**
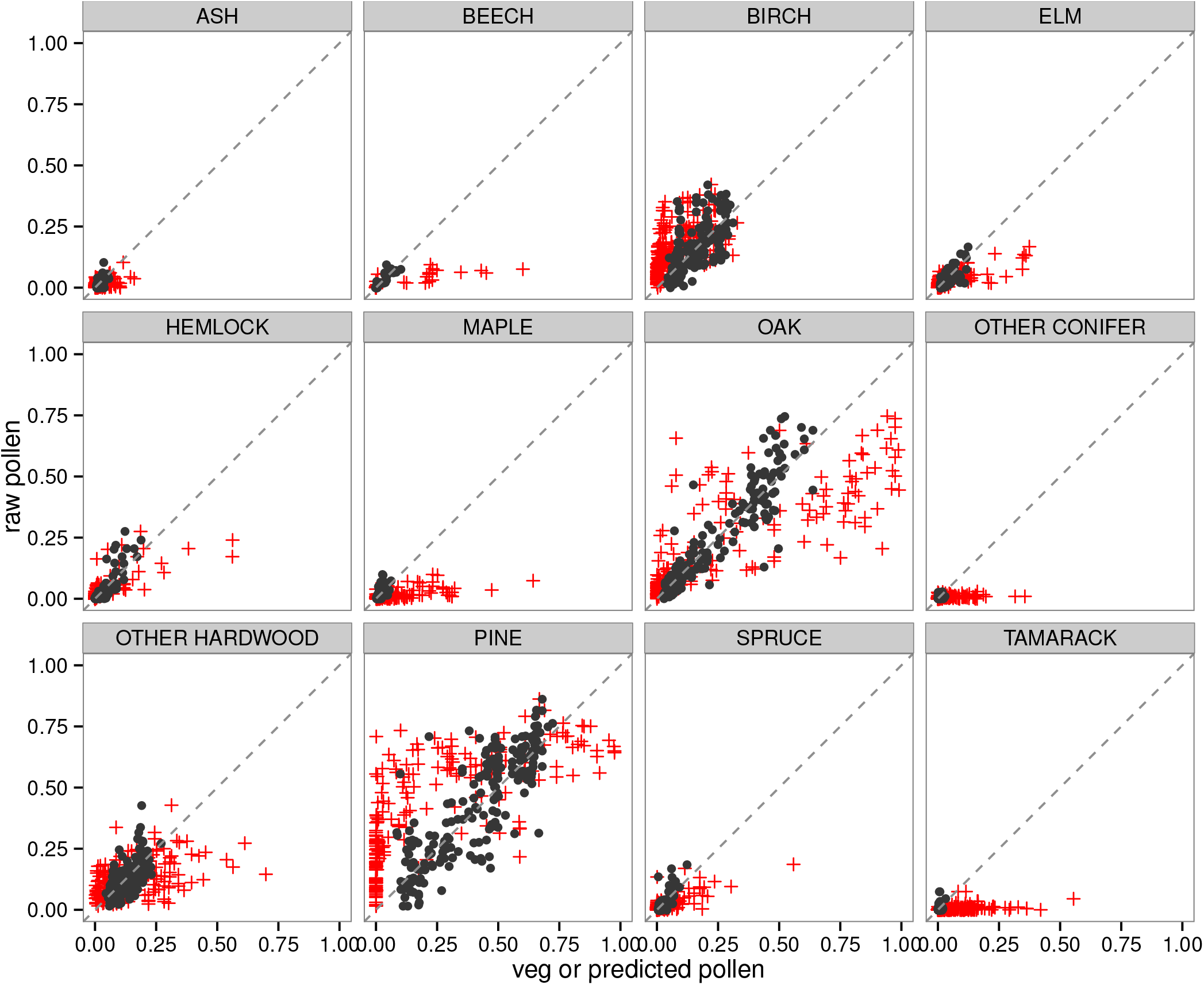
Pollen proportions plotted against local vegetation proportions (red crosses) or model-predicted pollen for the variable power-law kernel (PLK) model (black dots).

From each model we obtain taxon-specific estimates of pollen productivity *ϕ* (Fig. 5). If this parameter truly represents differential pollen production, then *ϕ* estimates should be more or less consistent across model variants, which is generally the case (Fig. 5). However, for most taxa, variable PLK model *ϕ* estimates are larger than for the other models (although perhaps not statistically different). The estimated values of *ϕ* form three distinct groups in *ϕ* parameter space: low, intermediate, and high production (Fig. 5). Taxa with lowest production in decreasing order are beech, maple, other conifer, and tamarack. This low production pattern is evident in Fig. 4, where the representation of these taxa in the pollen records is consistently less than their representation on the landscape (except at a few anomalous ponds). Intermediate producers, again in decreasing order, include other hardwood, oak, elm, hemlock, ash, and spruce. The high production group includes pine and birch, which can also be seen in Fig. 4 by their propensity to be over-represented in the pollen record relative to the landscape. These results agree in general with Prentice and Webb (1986), who analyzed the pollen-vegetation relationship for sites in Wisconsin and the Upper Peninsula of Michigan. In that work, pine and birch were identified as high producers/dispersers, while maple and tamarack were identified as low producers/dispersers. Jackson (1990) also found that pine and birch were good dispersers, while beech, maple, and tamarack had smaller source areas, indicating more-limited dispersal.

**Figure 5:**
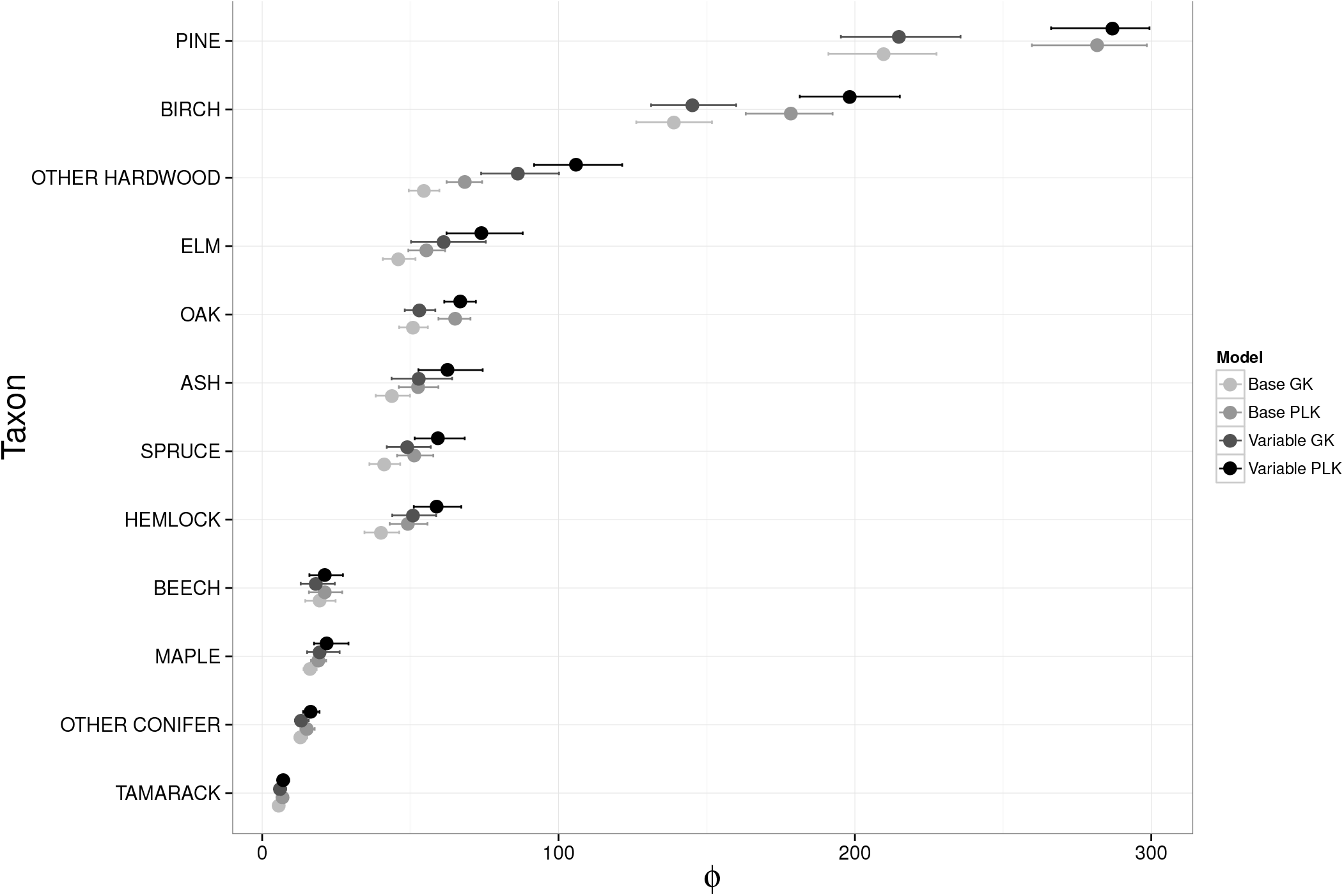
Mean values and 95% credible intervals for differential production parameter *ϕ*, by taxon for the four considered models (GK and PLK base and variable).

For the base models, the local versus non-local parameter *γ* had an estimated mean of 0.21 for the GK model and 0.046 for the PLK model. This indicates that roughly 21% (or 5%) of the pollen produced by vegetation in a focal grid cell is deposited in that grid cell, while the remaining 79% (or 95%) disperses elsewhere.

When we allowed *γ* to vary by taxon, the estimates for both kernels yielded similar results with overlapping 95% credible intervals for all taxa except pine and oak (App. A, Table A). For both kernels, birch, ash, other hardwood and other conifer all had lower estimates of *γ*, with mean posterior values all less than 0.10 (mean values ranging from 0.015 - 0.088). Pine and oak estimates of *γ* were also small for the variable PLK model (*γ* < 0.05), but not for the variable GK model (*γ* > 0.2). Both hemlock and tamarack had larger estimated values of *γ* for both models. It is encouraging that these estimates of taxon-specific local strengths are similar among models, but based on these estimates alone we can compare only focal-cell contributions. Non-local dispersal is determined by the product of (1 − *γ*) and the sum of the weights of all non-local cells, which in turn depends on the estimated kernel parameter(s).

To assess how *γ* and the kernel parameters affect pollen dispersal, we plot the proportion of deposited pollen as a function of distance from the pollen source. More precisely, for each taxon we plot the cumulative density function (CDF) given by *γ* + (1 − *γ*) Σ*_sj ≠ si_ w*(*s_i_,s_j_*) as a function of radius from the source, where cells are included in the non-local sum if the distance between *s_i_* and *s_j_* is less than or equal to this radius (Fig. 6). The weights *w*(*s_i_, s_j_*) depend on the estimated kernel parameters, whose quantities are not directly interpretable on their own but are reported in Appendix A, Table B. Note that the base model CDFs do not vary by taxon (Fig. 6, solid lines).

**Figure 6:**
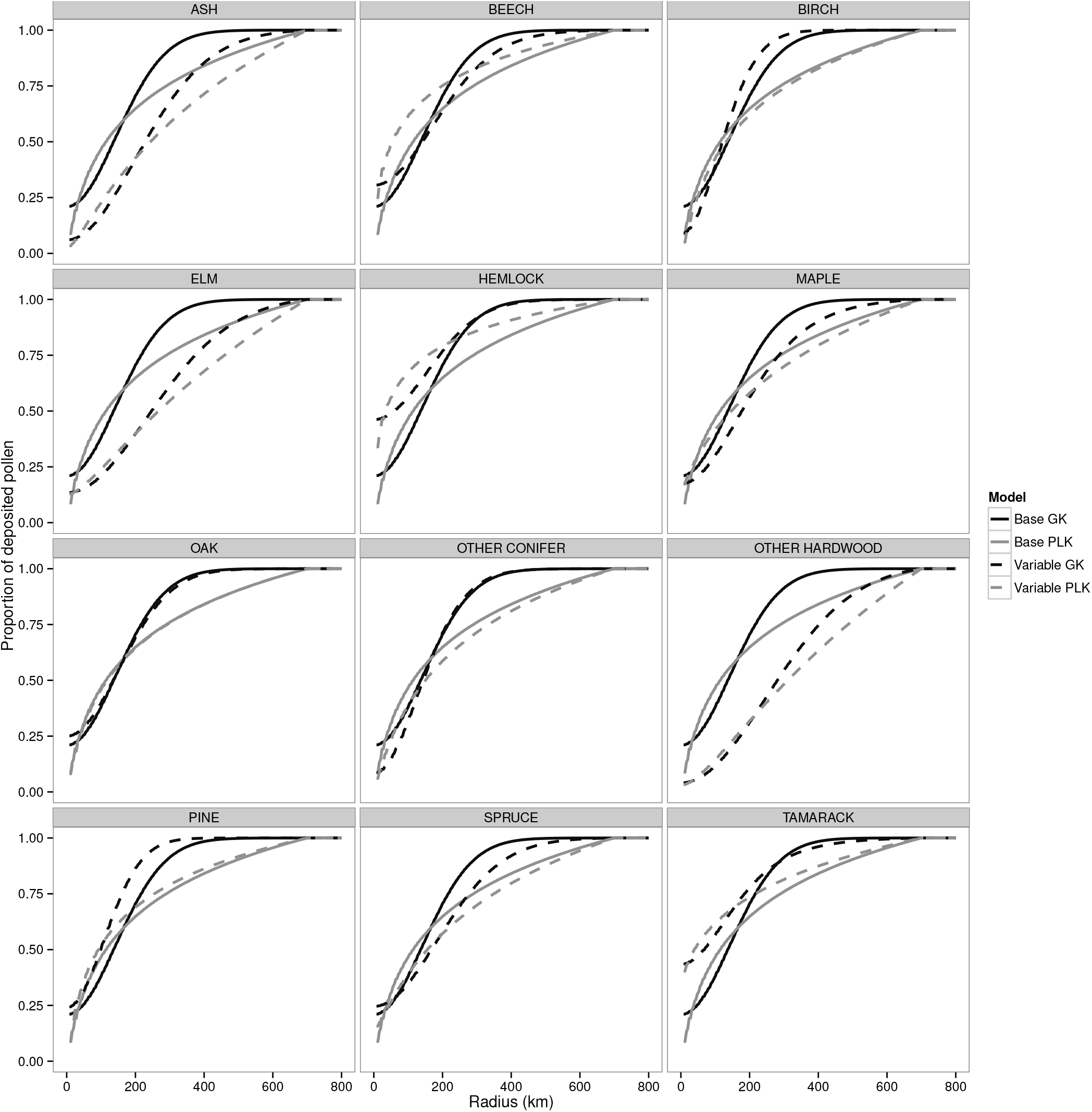
The proportion of deposited (or accumulated) pollen plotted as a function of radius for each of the modelled taxa and for each of the four considered models (GK and PLK base and variable).

As seen in Fig. 6, the base model CDFs have two points of intersection, indicating changing relative rates of pollen accumulation. CDFs for the GK models have a sigmoidal shape, where smaller values of *ψ* correspond with steeper slopes indicating that relatively more pollen is deposited close to the source. This effect is most obvious when we contrast the variable GK CDFs for pine (lowest *ψ*) and other hardwood (greatest *ψ*). For the variable PLK model, for fixed b, larger values of a result in a more sigmoidal shape that resembles the GK, as seen in the case of elm. Again for fixed *b*, smaller values of a result in *a* concave downward curve indicating more deposition closer to the source, but at a certain distance from the source - the radius at which the GK and PLK CDFs intersect - the cumulative pollen deposition is equal. To condense these results, we determined the radius from the source needed to capture 50, 70, and 90% of the deposited pollen for each model by taxon (Table 2). The 50% capture radius is smaller for the PLK base model than for the GK base model. However, the CDF for the GK base model asymptotes to one at a smaller radius than the PLK base model (GK base has a thinner tail). For the variable models, the 50% capture radius was smallest for hemlock (60 km for the variable GK; 32 km for the variable PLK) and largest for other hardwood (280 km for the variable GK; 312 km for the variable PLK). The 90% capture radii ranged from 216 km for pine to 512 km for other hardwood for the variable GK, and 380 km for elm to 608 km for other hardwood for the variable PLK. These values are larger than previous estimates of source area. Williams and Jackson (2003) estimated source area by comparing modern broad- and needle-leaved pollen percentages to forest cover obtained via remote sensing, and found that average source areas appear to be from 25 - 75 km. Prentice et al. (1987) and Bradshaw and Webb (1985) also estimated that for small to medium lakes the majority of the pollen comes from a source area on the order of 30 - 50 and 10 - 30 km respectively. Conversely, a modeling study found that 30% pollen-source radii ranged between 50 and 800 m for lakes < 20 hectares, and that vegetation sampling restricted to this area was sufficient to obtain reliable parameter estimates Sugita (1994). In subsequent work, however, Sugita (2007*b*) noted that for most taxa, 90 - 95% of pollen comes from within a source area of 100 - 400 km of the depositional basin.

**Table 2:** Radii (km) from a pollen source needed to capture 50, 70 and 90% of the dispersed pollen for the variable Gaussian and variable power-law models. The Gaussian base model estimated radii of 136 (128; 144), 200 (188; 208) and 292 (276; 308) km to capture 50, 70 and 90% of the dispersed pollen across all taxa, while the power-law model estimated radii of 116 (104; 124), 240 (228; 256) and 492 (484; 504) km. ^*^The Other hardwood grouping is composed of: Alnus, Carya, Cornus, Gleditsia, Liquidambar, Maclura, Morus, Nyssa, Ostrya carpinus, Platanus, Populus, Robinia, Rosaceae, Salix, Tilia, Zanthoxylum, as well as any undifferentiated hardwood (e.g. pollen grains classified as Fagus/Nyssa, Betula/Corylus, etc.).

| Taxon names |  | Model |  |  |  |  |  |
| --- | --- | --- | --- | --- | --- | --- | --- |
|  |  | Gaussian |  |  | Power-law |  |  |
|  |  | 50% | 70% | 90% | 50% | 70% | 90% |
| Common | Scientific |  |  |  |  |  |  |
| Hemlock | *Tsuga* | 60 ( 8, 104) | 168 (136, 196) | 284 (248, 328) | 32 ( 20, 56) | 116 ( 84, 164) | 380 (332, 436) |
| Tamarack | *Larix* | 68 ( 8, 196) | 156 ( 72, 344) | 268 (156, 532) | 40 ( 8, 188) | 152 ( 44, 372) | 428 (284, 580) |
| Pine | *Pinus* | 100 ( 88, 108) | 148 (136, 160) | 216 (200, 240) | 92 ( 80, 104) | 208 (188, 228) | 468 (440, 484) |
| Birch | *Betula* | 120 (108, 136) | 164 (148, 188) | 232 (204, 264) | 132 (104, 160) | 260 (224, 300) | 504 (476, 528) |
| Oak | *Quercus* | 136 (120, 152) | 204 (188, 224) | 304 (276, 336) | 116 (100, 140) | 244 (216, 272) | 492 (472, 512) |
| Beech | *Fagus* | 144 ( 8, 192) | 228 (180, 292) | 348 (280, 452) | 52 ( 24, 100) | 152 ( 96, 228) | 420 (352, 484) |
| Other conifer | *Abies* & *Cedrus* | 144 (112, 184) | 196 (152, 252) | 276 (216, 352) | 148 ( 96, 212) | 284 (216, 356) | 520 (472, 564) |
| Spruce | *Picea* | 168 (132, 212) | 252 (208, 316) | 372 (308, 464) | 156 (108, 220) | 300 (236, 372) | 532 (488, 576) |
| Maple | *Acer* | 180 (108, 320) | 256 (160, 432) | 368 (236, 580) | 144 ( 56, 324) | 288 (148, 468) | 524 (416, 620) |
| Ash | *Fraxinus* | 228 (176, 304) | 308 (236, 408) | 432 (332, 552) | 244 (168, 324) | 384 (304, 460) | 580 (532, 616) |
| Elm | *Ulmus* | 248 (192, 328) | 344 (268, 444) | 484 (384, 588) | 268 (196, 344) | 416 (344, 484) | 596 (556, 628) |
| Other hardwood | * | 280 (240, 328) | 376 (320, 432) | 512 (444, 572) | 312 (260, 360) | 448 (400, 488) | 608 (588, 628) |

To visualize the predicted dispersal patterns, we estimate the proportion of deposited pollen by taxon for each grid cell in the domain (Fig. 2; previously, as seen in Figs. 3 and 4, we estimated deposited pollen only for grid cells with pollen data). These spatial maps illustrate how the models treat dispersal - the predicted pollen maps for the variable PLK model appear to be more smooth. This difference in part attributed to the shallow slope of the PLK relative to that of the GK; more pollen is deposited at short/intermediate distances, resulting in a gentle gradient (see Fig. 6).

We compare predicted pollen maps with maps of the PLS data (Fig. 2), recalling that pollen predictions have been scaled to account for differential production (i.e., we compare predicted pollen with vegetation composition). Accounting for differential production highlights the under-production of maple pollen; maple pollen proportions are consistently lower than vegetation proportions. Fig. 2 also illustrate the differences in pollen and vegetation ranges. The effects of dispersal are especially clear in the cases of pine and birch, where pollen disperses well beyond the vegetation range limits.

### 3.4 Assessing productivity and dispersal estimates

We compared the STEPPS variable PLK model productivity estimates, *ϕ*, to sets of published PPEs (Fig. 7). PPEs included in the comparison were slope coefficients in either a linear regression, ERV, or bias-corrected ERV model. PPEs were also split into geographical regions based on the sampling locations, which were upper Midwestern USA (UMW), northeastern USA, and the European continent.

**Figure 7:**
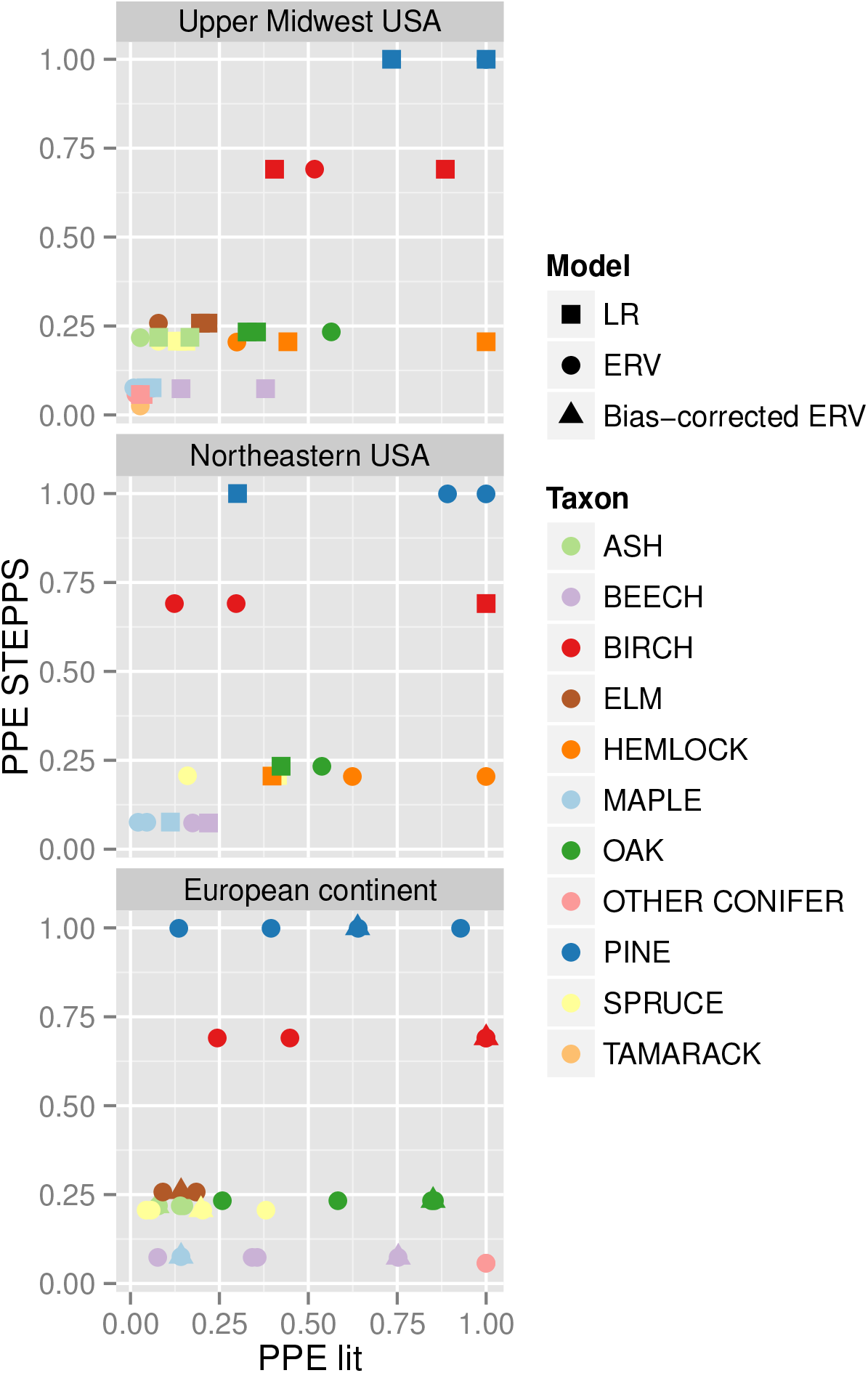
Pollen productivity estimates (PPEs) from STEPPS versus sets of previously published PPEs from different geographical regions (Upper Midwest USA, Northeastern USA, European continent). Published PPE data sets were generated using linear regression (LR), extended R-value (ERV), or bias-corrected ERV models.

STEPPS productivity estimates are most similar to PPEs from the UMW in terms of relative rank, while PPEs further from the UMW agree less with the STEPPS productivity estimates/rankings. However, a positive linear relationship exists between STEPPS productivity estimates and the PPEs from the northeastern USA. For the PPEs from the European continent the relationship is weak, although both STEPPS and PPE estimates agree that ash, elm, maple, and spruce are low producers. The variability in PPEs across space and the closer agreement of STEPPS productivity estimates with PPEs from the UMW is likely a result of the species composition and forest structure of a particular region. The strength of the relationship between STEPPS estimates and PPEs is not strongly affected by the method used to estimate the PPEs.

Next, we compared our estimated 90% capture radius, which we defined as the radius of the circle (centered at the pollen source) within which 90% of the pollen is deposited, to published values of falling speed for trees from continental Europe and North America within the same genus as the STEPPS taxa (Fig. 8). Capture radius is similar to the widely used concept of relevant pollen source area (Sugita, 1994), but calculated from point of source rather than point of deposition. Falling speed values used in this comparison were obtained from the compilation of Jackson and Lyford (1999).

**Figure 8:**
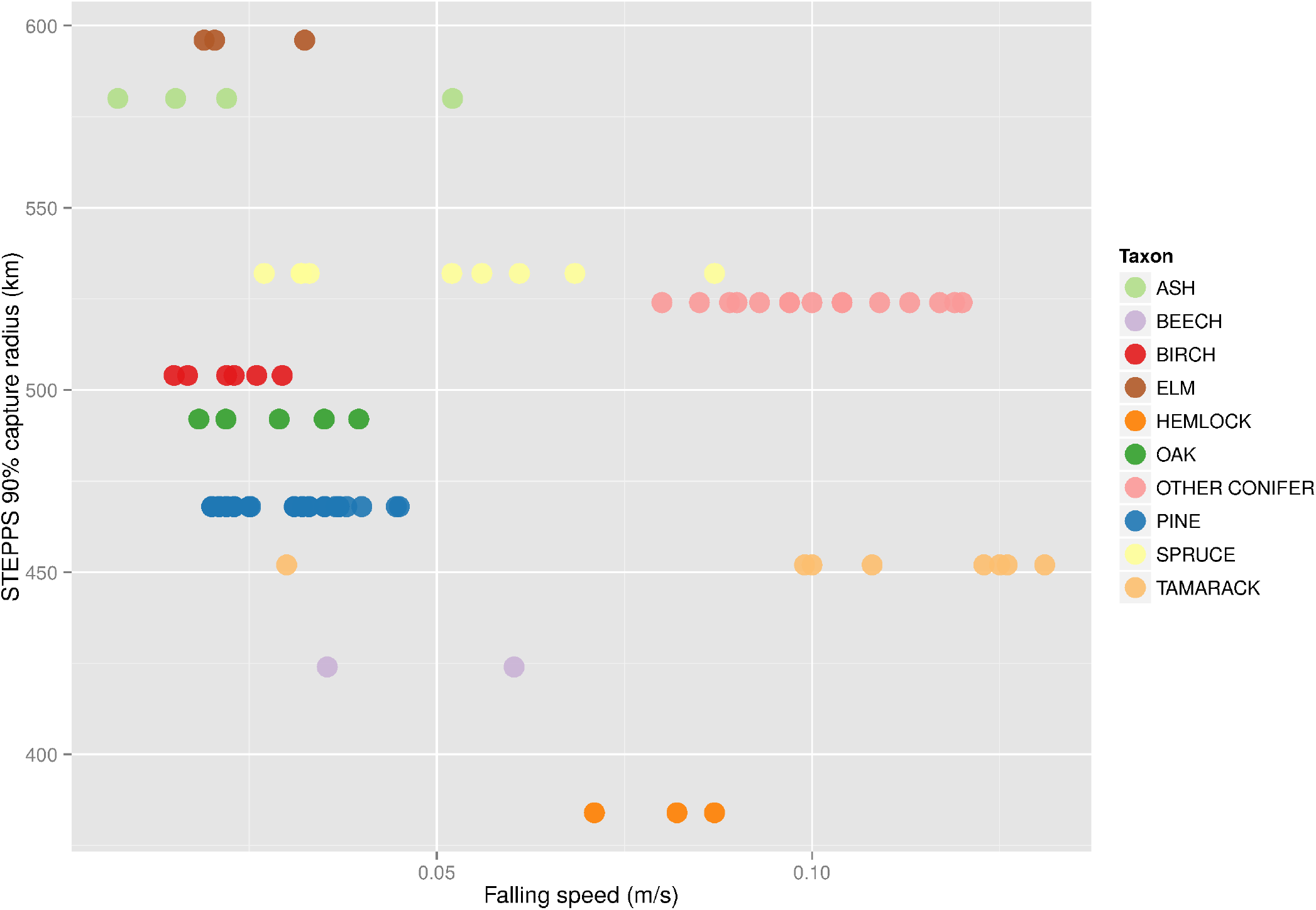
Scatter plot of the STEPPS 90% capture radius versus measured falling speeds.

As expected, the STEPPS capture radius and falling speed are inversely related; a lower falling speed corresponds with a greater dispersal ability. Several taxa, however, do not fit this relationship: hemlock, maple, and spruce. Measured falling speeds vary for these and other taxa, both within and among laboratories and methods (Jackson and Lyford, 1999).

These comparisons provide additional support for the STEPPS model; we find agreement between STEPPS estimates and our current understanding of production and dispersal processes.

### 3.5 Vegetation composition predictions

We estimated settlement-era vegetation using the prediction model with four candidate calibration models (GK and PLK, base and variable). The model was fit using a modified version of Stan with a warm-up period of 150 iterations and a sampling period of 1000 iterations. Parameter effective sample sizes were small (10-200) reflecting the complex dependence structure of the model. However, trace plots indicated successful mixing.

Predicted patterns of settlement-era composition are generally similar across models and dissimilarities among maps are low (App. A, Figs. B & C). According to the Euclidean distance, composition estimates with the underlying base PLK calibration model are closest to the data (Table 1, Distance = 1979), and farthest for the GK calibration model (Distance = 2078). For most taxa, the STEPPS composition predictions (using, for illustration purposes, the variable PLK) broadly match the compositional distributions in the PLS data (Fig. 9). As expected, the STEPPS predictions appear smoother; this is a result of the long-range pollen transport smoothing out local features, but also because of the limited number of pollen sample sites. Smaller, compositionally unique regions such as the Minnesota Big Woods are not well resolved by the model (App. A, Fig. B).

**Figure 9:**
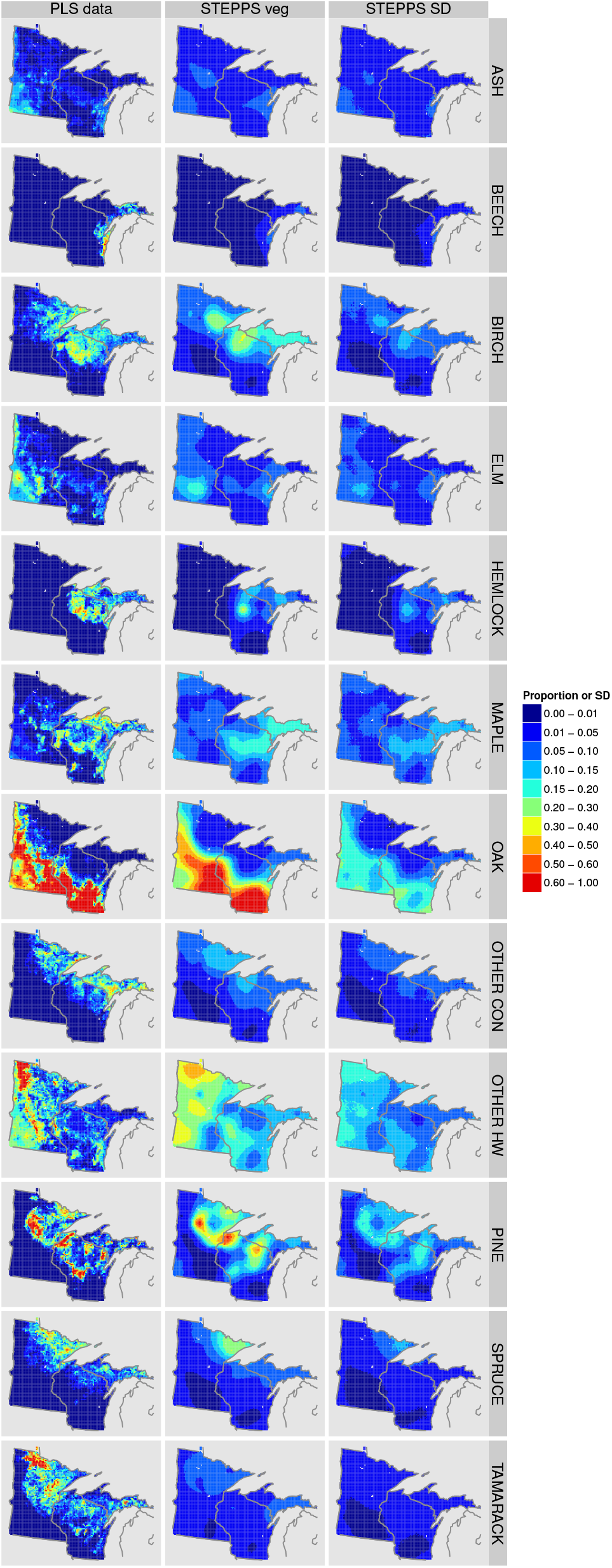
Heat maps of the PLS data, STEPPS composition estimates and the standard deviation of the posterior sample, based on the variable PLK model.

STEPPS does not adequately predict the spatial distribution of tamarack; in Wisconsin tamarack is highly underpredicted (App. A, Fig. B), and dispersal credible intervals indicate large uncertainties (Table 2). Tamarack is a low pollen producer and a poor disperser (Janssen, 1966; Webb et al., 1981). Populations are highly concentrated in localized settings, such as peatlands, but are rare across the uplands. Given this, we expect to observe tamarack pollen only in peatlands or lakes in peatland-rich areas, and even there the tamarack signal will usually be swamped by pollen from better producers and/or dispersers. Since there are few pollen records from tamarack-dominated locations, it is not surprising that the predictive ability of STEPPS is poor for tamarack.

## 4 Discussion

Bayesian pollen-vegetation models (PVMs) enable spatial reconstructions of vegetation composition and estimation of key process parameters, while accounting for data, process, and parameter uncertainty, as well as complex spatial dependencies (Paciorek and McLachlan, 2009). Here we work towards improving reconstructions of vegetation and quantifying their uncertainty using a Bayesian hierarchical PVM (STEPPS) calibrated with newly developed settlement-era vegetation and pollen datasets. STEPPS is not derived from physical first principles, but it accounts for differential pollen production and dispersal among taxa. The STEPPS PVM is able to estimate pollen-vegetation process parameters without the need to quantify any taxon-specific attributes *a priori*; these estimates are collectively informed by the network of pollen sites and PLS data. This is in contrast to some other PVMs, such as LOVE and REVEALS (Sugita, 2007*a,b*), which require prescribed estimates of taxon-specific fall speeds and productivity, in addition to assumptions about local atmospheric conditions (Jackson and Lyford, 1999). Despite these differences, both PVMs have demonstrated their ability to predict vegetation composition (Soepboer et al., 2010; Wang and Herzschuh, 2011; Overballe-Petersen et al., 2013). Continued model comparisons and development will lead to better understanding of physical and ecological assumptions and parameters.

In STEPPS, we do not account for the effects of sedimentary basin size. Previous studies have found that basin size is related to pollen source area - the area from which deposited pollen originates (Jacobson and Bradshaw, 1981; Prentice, 1985; Sugita, 1994). Larger basins are less sensitive to local vegetation, and provide regional integration. As a result, pollen records from larger lakes are more frequently used to reconstruct regional vegetation. However, even the smallest basins (Jackson, 1990) and forest-floor pollen assemblages (Jackson and Kearsley, 1998; Calcote, 1995) have a strong regional signal. Recent work has shown that regional vegetation can be inferred from smaller-sized lakes (Trondman et al., 2015). In STEPPS we use all suitable pollen records to reconstruct regional vegetation. Vegetation composition is assumed constant within grid cells, and pollen dispersal to a basin is based on the distance from the cell centroid to the basin. The coarseness of the PLS data prevents resolution of sub-grid-cell heterogeneity in forest composition; as such, accounting for lake size would likely not improve predictions. However, the STEPPS PVM can be refined to account for and assess the importance of basin size.

Following standard convention in Quaternary palynology, we assume that dispersal is isotropic, and that it does not vary throughout the domain. In reality, pollen dispersal is a complex function of many factors, including wind direction and speed and topographical and landscape features. Asymmetric dispersal has been documented by Robledo-Arnuncio and Gil (2005) using DNA markers to identify parentage, and non-uniform wind fields have been incorporated into a few PVMs (Bunting and Middleton, 2005). However, pollen dispersal is an underexplored problem and the importance of anisotropy has yet to be determined with any certainty. Frequent changes in wind direction and the multiple mechanisms and processes affecting pollen transport that act at different spatial and temporal scales suggest that a simple characterization of anisotropic dispersal may not improve predictions. It is possible to account for such dispersal-related complexities in a PVM calibrated during modern times for which wind data is available. However, covariate data needed to determine non-uniform dispersal patterns, such as wind speed, turbulence, and direction, are not readily available for past times. Regardless, further investigation is warranted.

Even with the simplifying assumption of isotropy, dispersal-kernel shape can vary. In these analyses, we used the GK and PLK as representative of short- and long-tailed kernel types. Predictions based on these kernels are similar, although the fatter-tailed PLK performed slightly better than the GK. This agrees with findings by others, who found evidence supporting the importance of long-distance pollen dispersal (Austerlitz et al., 2004; MacInnis, 2012), although additional data and analyses are needed to support these conclusions.

Most PVMs are calibrated with modern vegetation and pollen data, and are subsequently used to reconstruct past vegetation under the assumption that the pollen-vegetation relationship is constant. However, Euro-American settlement resulted in significant changes to the landscape that may have influenced the pollen-vegetation relationship, and calibration based on settlement-era data may improve past predictions (Kujawa et al., 2016; St Jacques et al., 2014). That being said, PVM calibration using modern data is still valuable; in many cases no past forest data are available, and in all cases, spatial precision and density of vegetation sampling is inevitably lower. Also, if it exists, prediction bias can likely be quantified and corrected for. In future work we will reconstruct vegetation using the STEPPS PVM calibrated with modern data and compare with settlement-era-calibrated predictions.

PVMs calibrated with modern data generally do not rely on age-depth models; modern pollen samples are taken from the upper-most sediment. However, past vegetation reconstructions based on dated pollen samples assume sediment age is known. Age-depth models are used to infer sediment age, and are based on chronological controls including radiometric and biostratigraphic dates. However, confidently estimating placement of biostratigraphic events including the Euro-American settlement horizon remains an underappreciated source of uncertainty and potential bias in paleoecology. Soliciting expert opinion has become an increasingly common way to assign parameter priors in Bayesian statistics (Choy et al., 2009). Here, we used expert elicitation to identify the settlement horizon in pollen records to reduce expert bias. Experts were in complete agreement for only about a third of the pollen records, indicating substantial uncertainty in identifying biostratigraphic signals. Further investigation can determine the effects of this uncertainty on PVM calibration and reconstructions; here we use elicitation only to reduce bias in identifying the settlement horizon.

The STEPPS PVM outperforms vegetation predictions based on raw pollen counts alone, as well as those based simply on scaling using pollen productivity (Figs. 3 and 4). Pine and birch have the highest pollen productivity estimates, consistent with prior productivity estimates (Bradshaw and Webb, 1985; Prentice and Webb, 1986). For the STEPPS estimates, long-distance dispersal plays a major role in determining pollen assemblages in the UMW domain. For some taxa, such as pine and birch, the variable PLK model estimates that 2% of the dispersed pollen remains within an 8 km grid cell, while the remainder is dispersed elsewhere. For other taxa, such as hemlock, 27% of the pollen remains local.

Our pollen capture-radius estimates are large compared to previous estimates (Sugita, 2007b, 1994; Williams and Jackson, 2003; Prentice et al., 1987; Bradshaw and Webb, 1985); capture-radii required to account for 90% of the pollen are on the order of hundreds of kilometres. A comparison of the PLS and pollen data (Fig. 1) shows evidence of long-distance-dispersal over large distances (e.g., high pine pollen abundances far outside the area of highest pine tree abundances). Differences between STEPPS and previous source area estimates may be in part a consequence of the 8 km spatial resolution of our vegetation data, which does not allow us to resolve within-grid-cell heterogeneity and the effects of local populations near depositional environments (Jacobson and Bradshaw, 1981; Bradshaw and Webb, 1985; Jackson, 1990). If local populations not evident in the PLS data contribute large amounts of pollen to depositional environments, the model will infer that this pollen originated from further afield. The importance of spatial resolution on source area estimates has been previously noted (Sugita, 1994). Multi-scale analyses that test the impacts of spatial resolution on pollen source area are needed to provide further insight.

The STEPPS PVM reproduces regional patterns of settlement vegetation for most taxa, with some smoothing attributed to long-distance-dispersal, model assumptions, mixing of sediments and air, and the limited number of pollen records (Fig. 2). Tamarack is the sole taxon not well-modeled by STEPPS; this poor performance is attributable to the low pollen production, poor pollen dispersal, and highly localized distribution of tamarack.

Overall, our application of STEPPS has demonstrated its ability to reconstruct regional patterns of vegetation in the UMW from a network of fossil pollen records, while providing new insights into the key processes governing pollen-vegetation relationships. Of course, in many cases vegetation composition data that are as extensive, complete, and rich as the PLS data is not available, although this is changing as hyperspectral remote sensing and LIDAR increasingly enable species-level forest mapping (Asner et al., 2012). Nevertheless, the modelling framework represented by STEPPS is useful for any region with independent spatial datasets of pollen assemblages and forest composition. Our ability to predict vegetation at pre-settlement era indicates that it should be possible to infer past forest composition robustly from fossil pollen data, and thereby better understand past forest dynamics during the changing climates and land use of the Holocene and Anthropocene.

## Acknowledgements

We thank Jeannine-Marie St. Jacques for participating in the elicitation exercise and for comments in the intellectual development stages of this work. We also thank paleoecologists who made their data available through the Neotoma Paleoecology Database or otherwise, especially Randy Calcote, Sara Hotchkiss, Beth Lynch, and Edward Cushing. We owe special thanks to Eric Grimm for his role in data and knowledge stewardship. For the PLS data, we thank David Mladenoff and others involved. Comments by Simon Brewer and an anonymous reviewer improved the paper. This work was carried out by the PalEON Project with support from the National Science Foundation MacroSystems Program through grants EF-1065702, EF-1065656, DEB-1241874, DEB-1241851 and DEB-1241868 and from the Notre Dame Environmental Change Initiative.

## Appendix A

**Table A:**
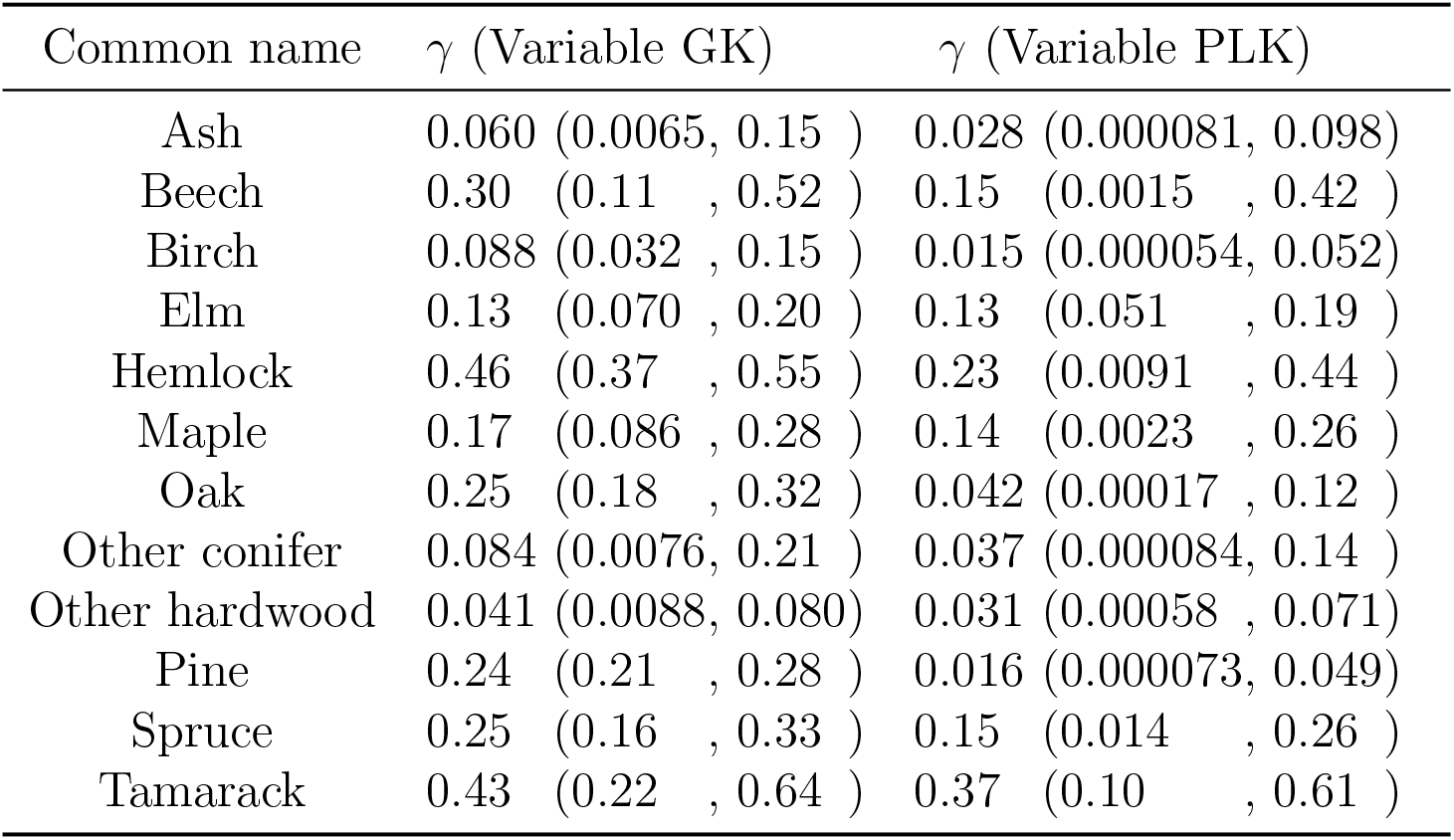
Estimates of parameter *γ*, indicating the proportion of locally sourced pollen for the variable Gaussian kernel (GK) model in which *ψ* and *γ* both vary by taxon and the variable power-law kernel (PLK) model in which a and *γ* vary by taxon. For the base models, *γ* was estimated to be 0.21 (0.19, 0.23) for the GK model and 0.046 (0.013, 0.078) for the PLK model.

| Common name | $\gamma$ (Variable GK) | $\gamma$ (Variable PLK) |
| --- | --- | --- |
| Ash | 0.060 (0.0065, 0.15 ) | 0.028 (0.000081, 0.098) |
| Beech | 0.30 (0.11 , 0.52 ) | 0.15 (0.0015 , 0.42 ) |
| Birch | 0.088 (0.032 , 0.15 ) | 0.015 (0.000054, 0.052) |
| Elm | 0.13 (0.070 , 0.20 ) | 0.13 (0.051 , 0.19 ) |
| Hemlock | 0.46 (0.37 , 0.55 ) | 0.23 (0.0091 , 0.44 ) |
| Maple | 0.17 (0.086 , 0.28 ) | 0.14 (0.0023 , 0.26 ) |
| Oak | 0.25 (0.18 , 0.32 ) | 0.042 (0.00017 , 0.12 ) |
| Other conifer | 0.084 (0.0076, 0.21 ) | 0.037 (0.000084, 0.14 ) |
| Other hardwood | 0.041 (0.0088, 0.080) | 0.031 (0.00058 , 0.071) |
| Pine | 0.24 (0.21 , 0.28 ) | 0.016 (0.000073, 0.049) |
| Spruce | 0.25 (0.16 , 0.33 ) | 0.15 (0.014 , 0.26 ) |
| Tamarack | 0.43 (0.22 , 0.64 ) | 0.37 (0.10 , 0.61 ) |

**Table B:**
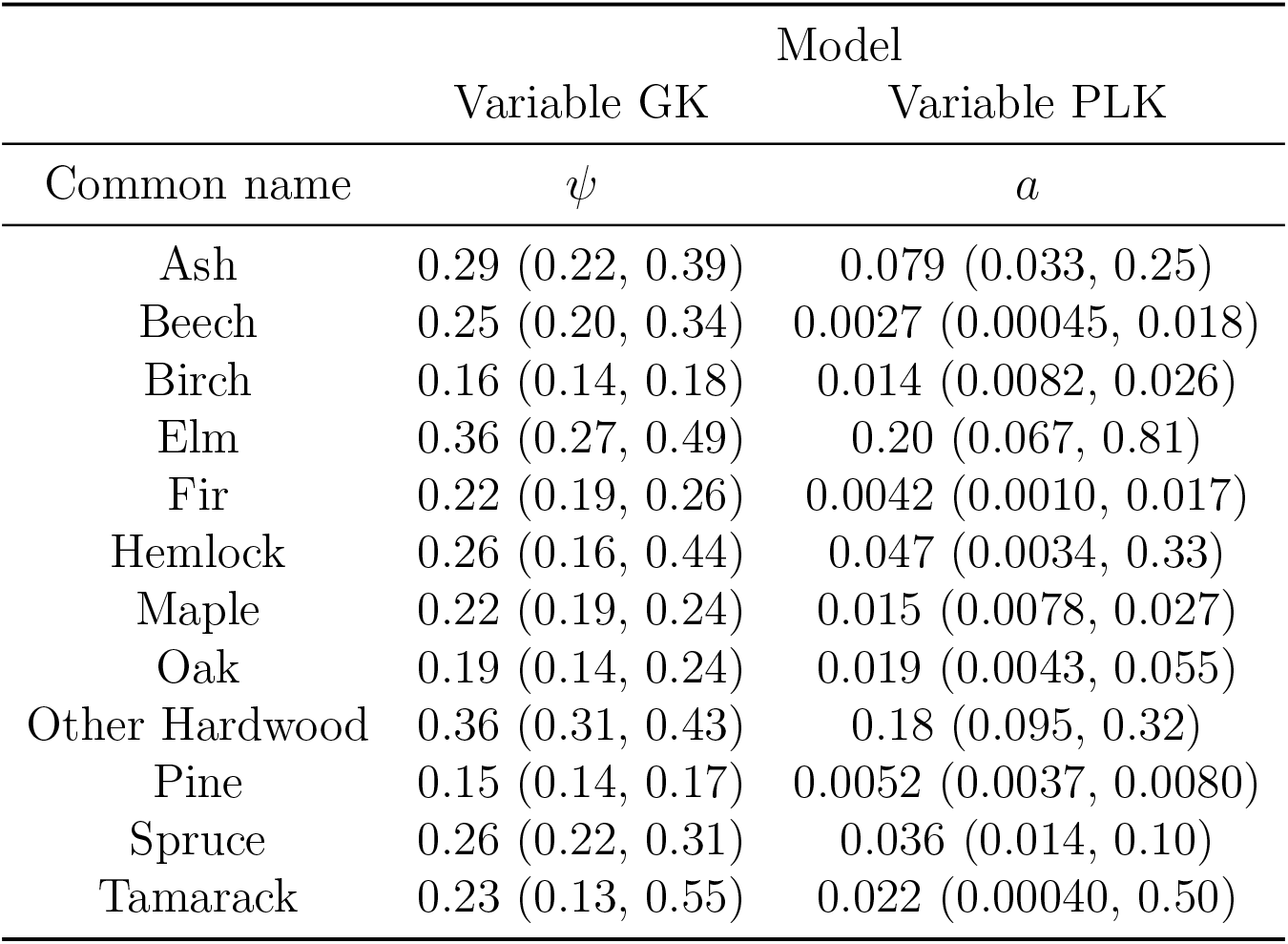
Estimated mean and 95% credible intervals for the taxon-specific dispersal kernel parameters for the variable Gaussian kernel (GK) and power-law kernel (PLK) models.

**Table C:**
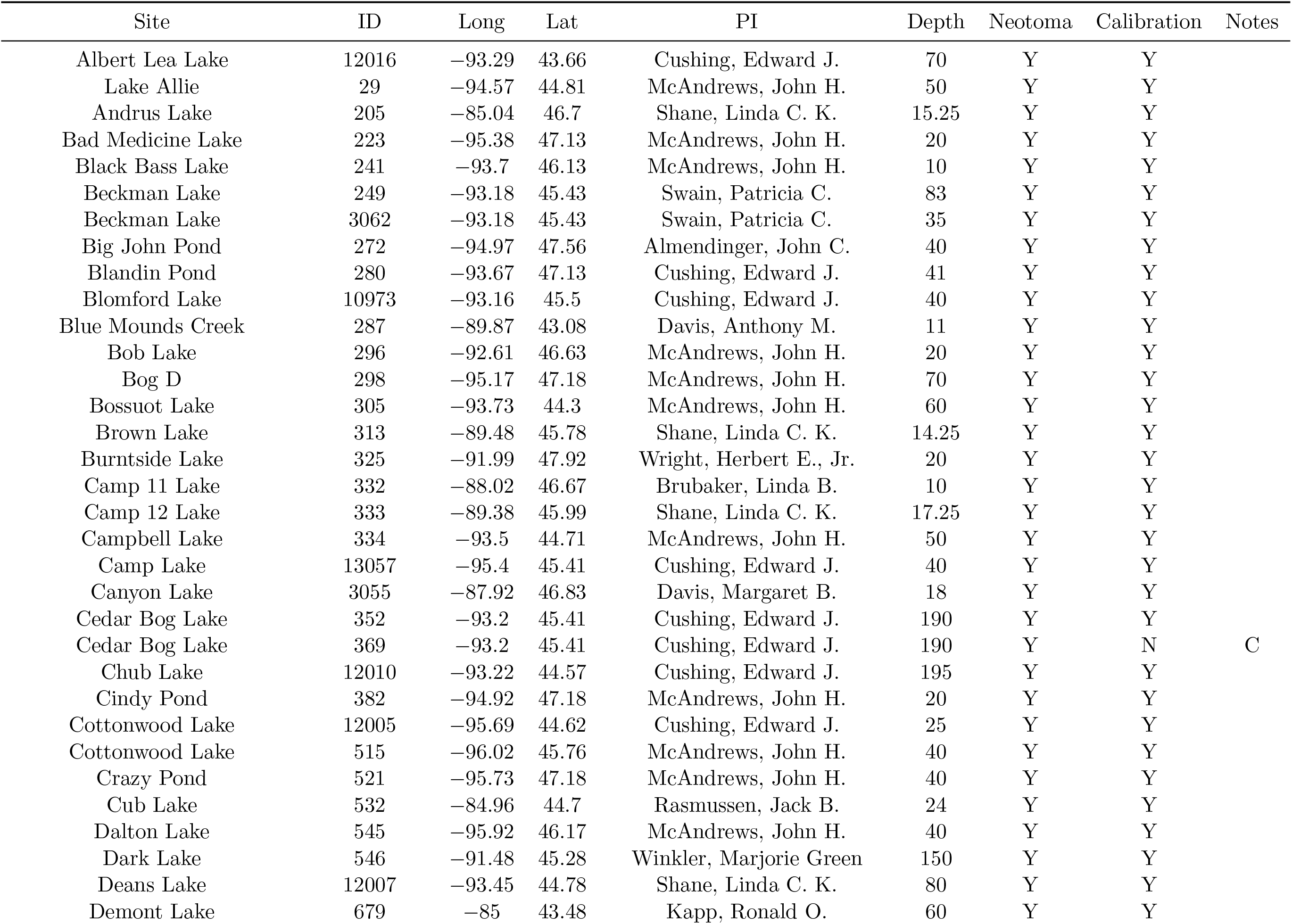
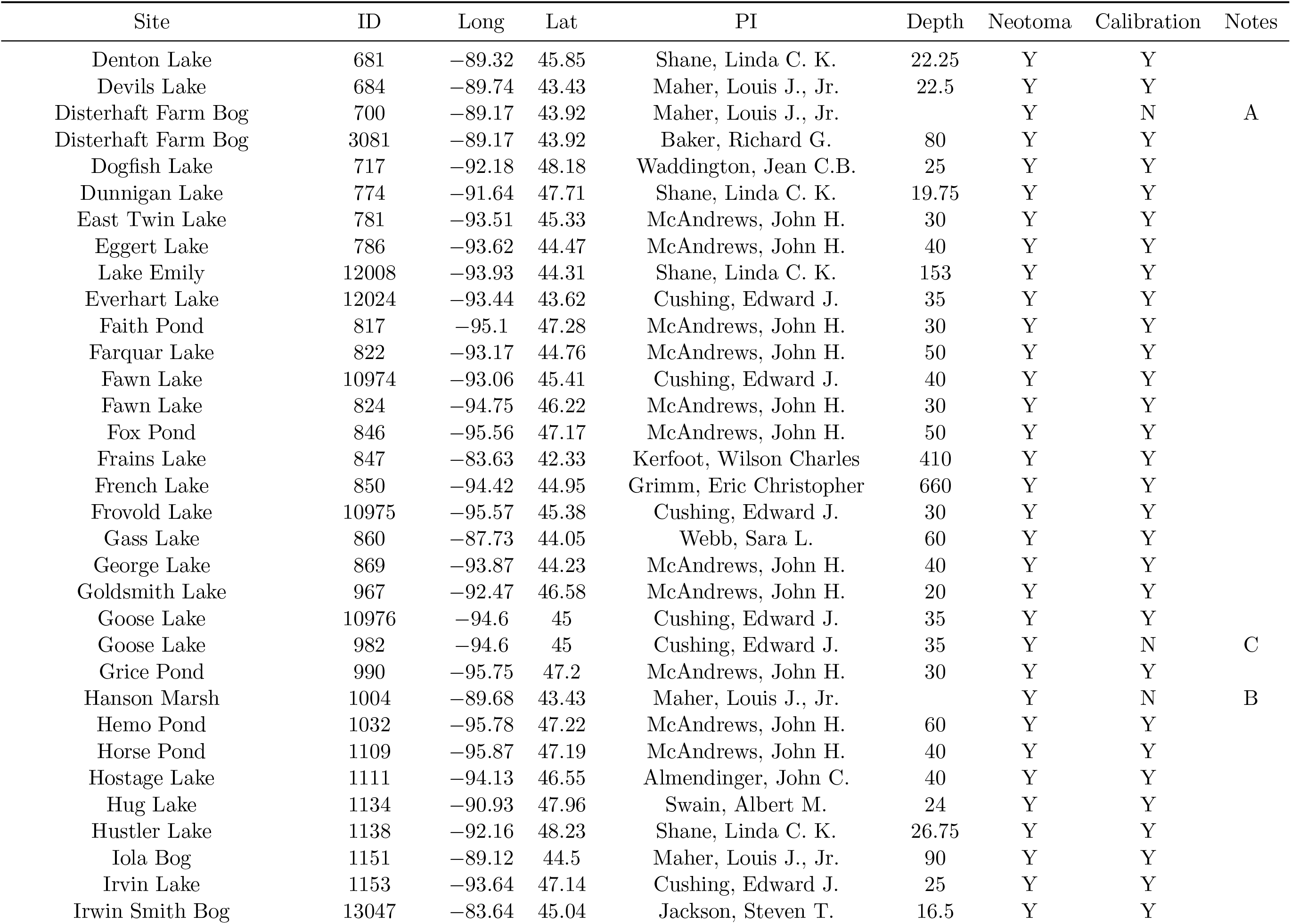
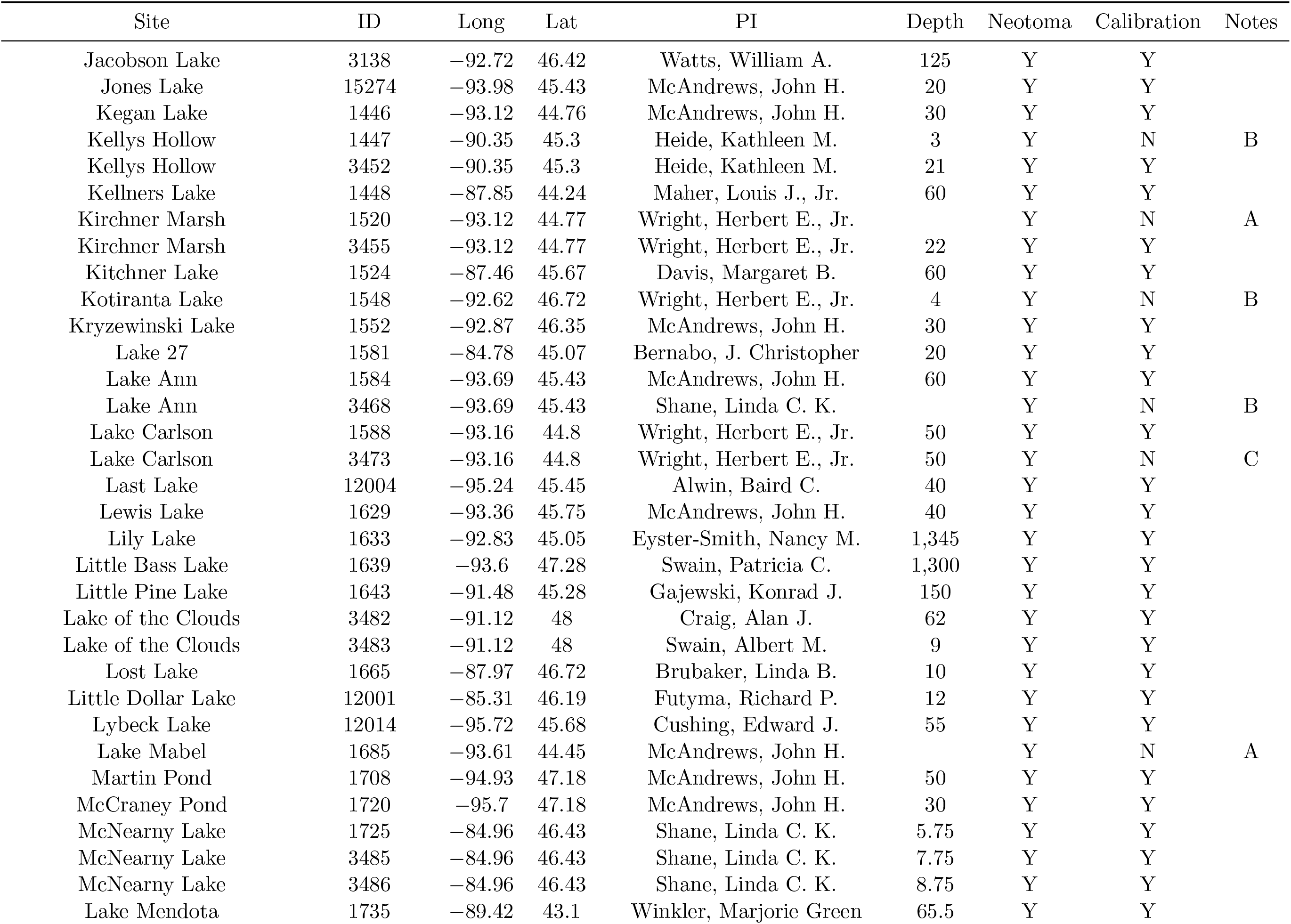
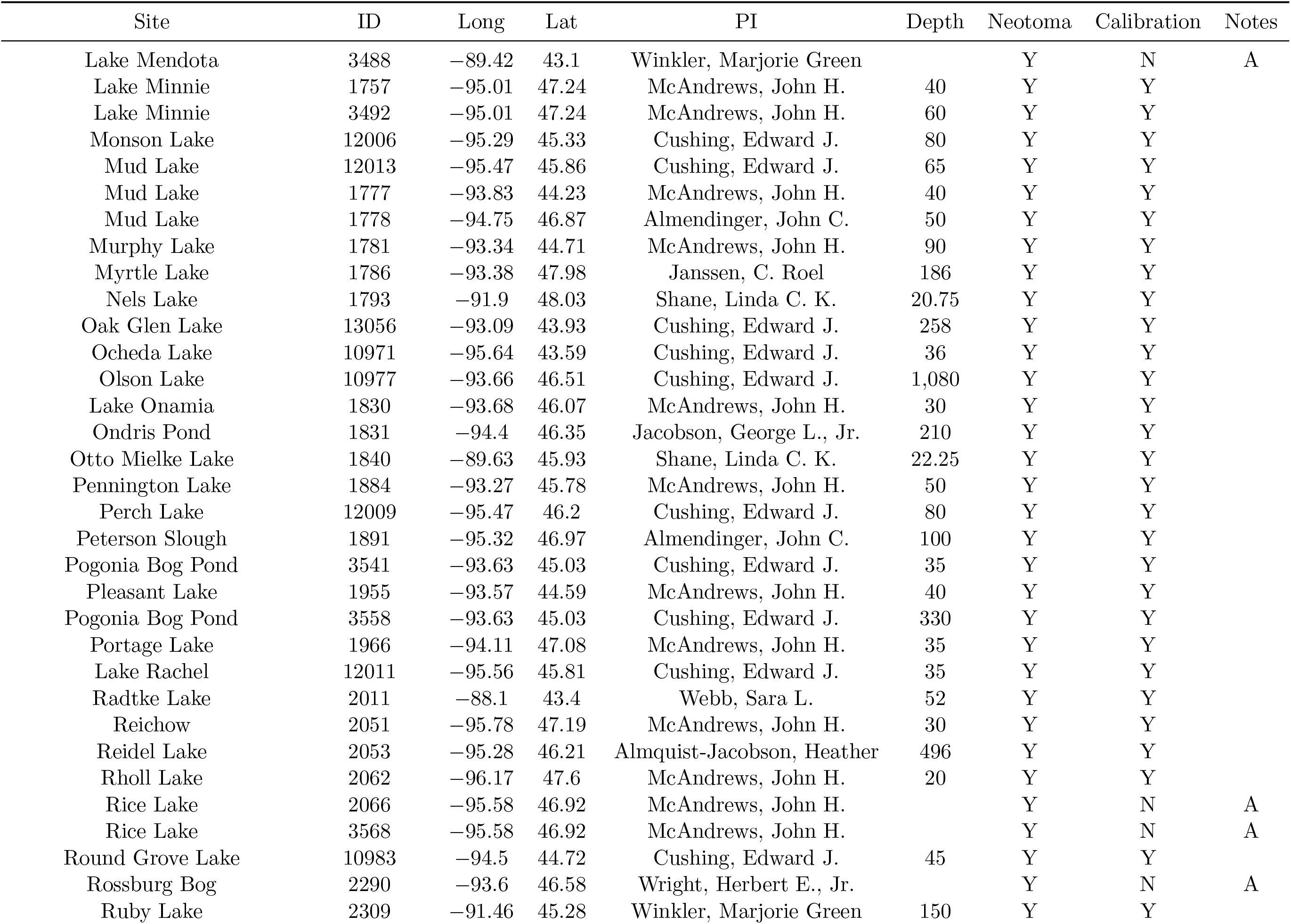
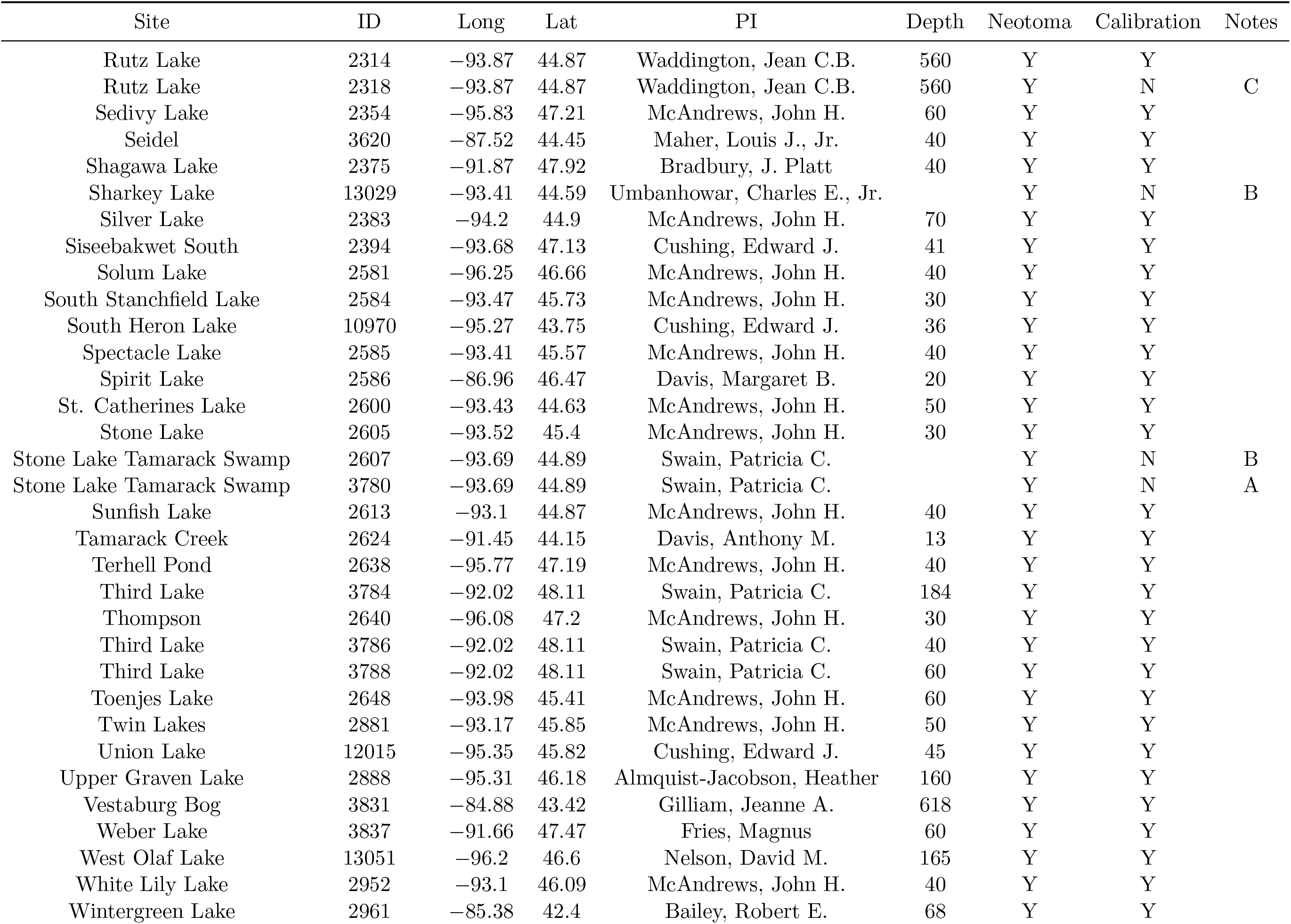
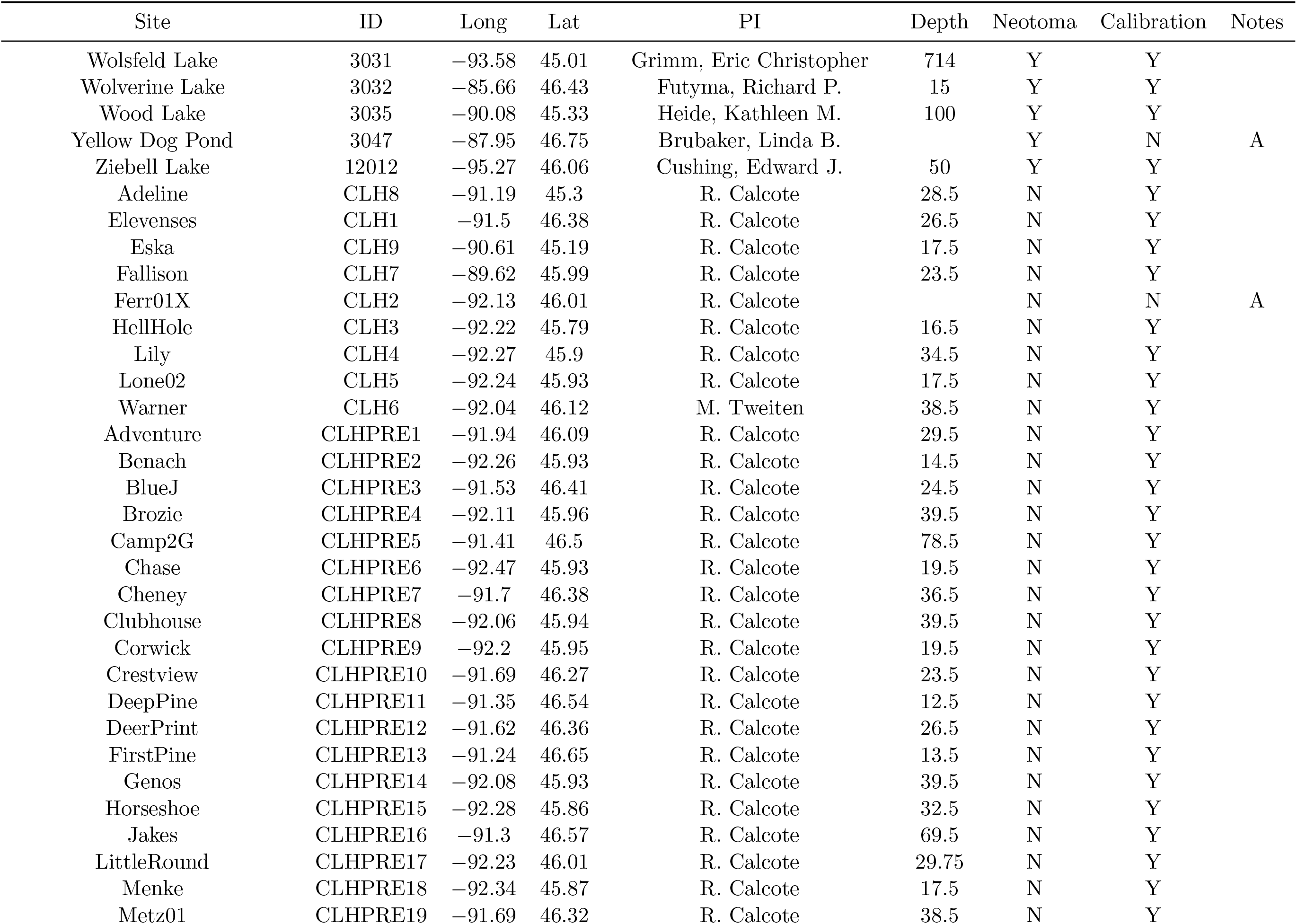
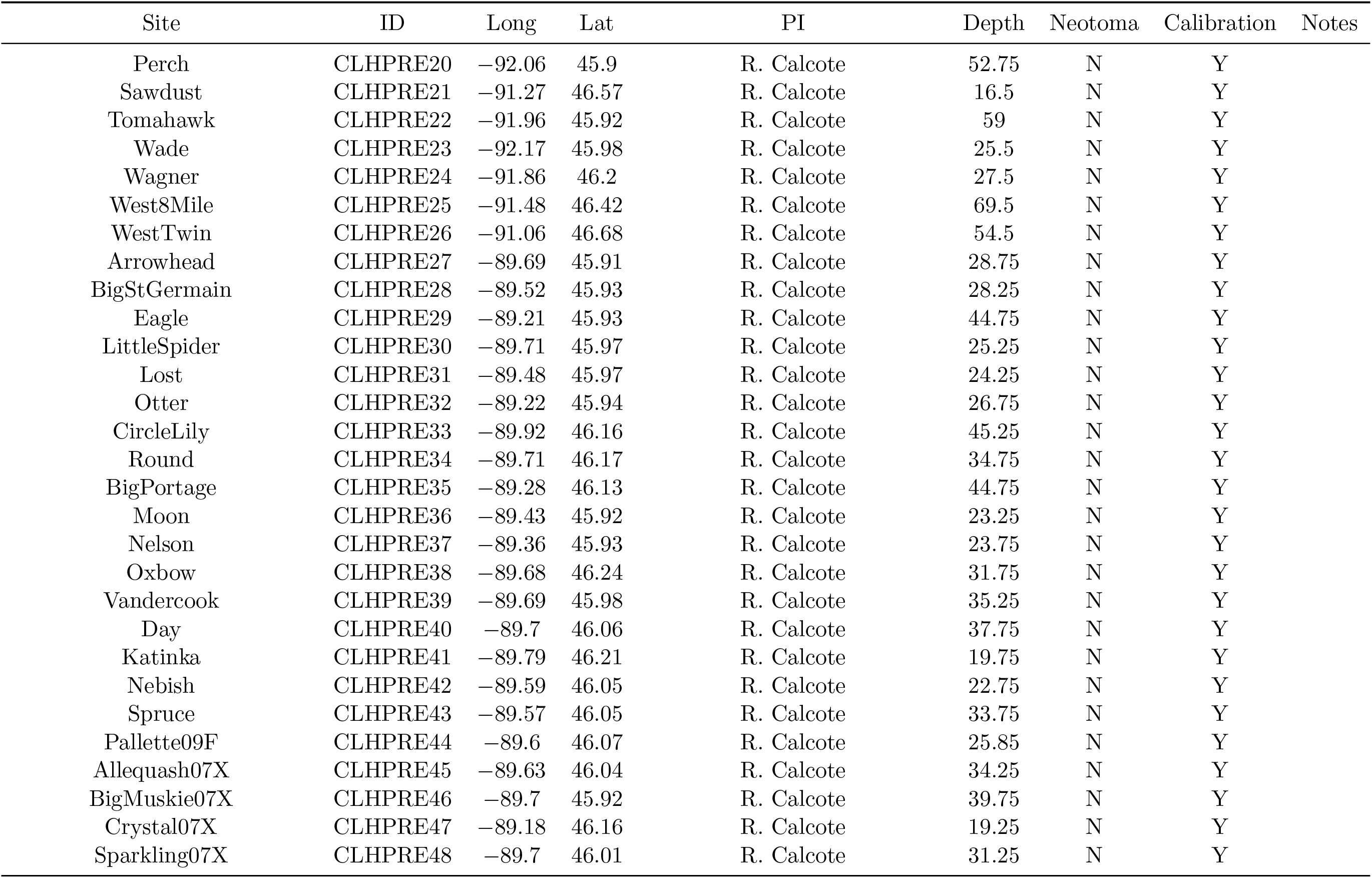
The sediment pollen sites in the upper Midwestern USA considered for the calibration data set. Metadata provided includes site name (Site), Neotoma or given identification number (ID), latitude (Lat) and longitude (Long), principal investigator (PI), settlement horizon sample depth (Depth; may be recorded as depth from the upper most sediment layer or from lake surface), an indication if the dataset came from Neotoma (Neotoma), an indication as to whether the dataset is included in the final calibration data set (Calibration), and in the case a site is not included (Calibration value of N), then a note indicating the reason for inadmissibility (Notes). Reasons for inadmissibility are: A) Three or four experts did not assign a pre-settlement sample; B) Two experts did not assign a settlement horizon sample, and the two assigned settlement horizon samples were at least 500 years away from 1850 according to the default age-depth models; C) No core top.

**Figure A:**
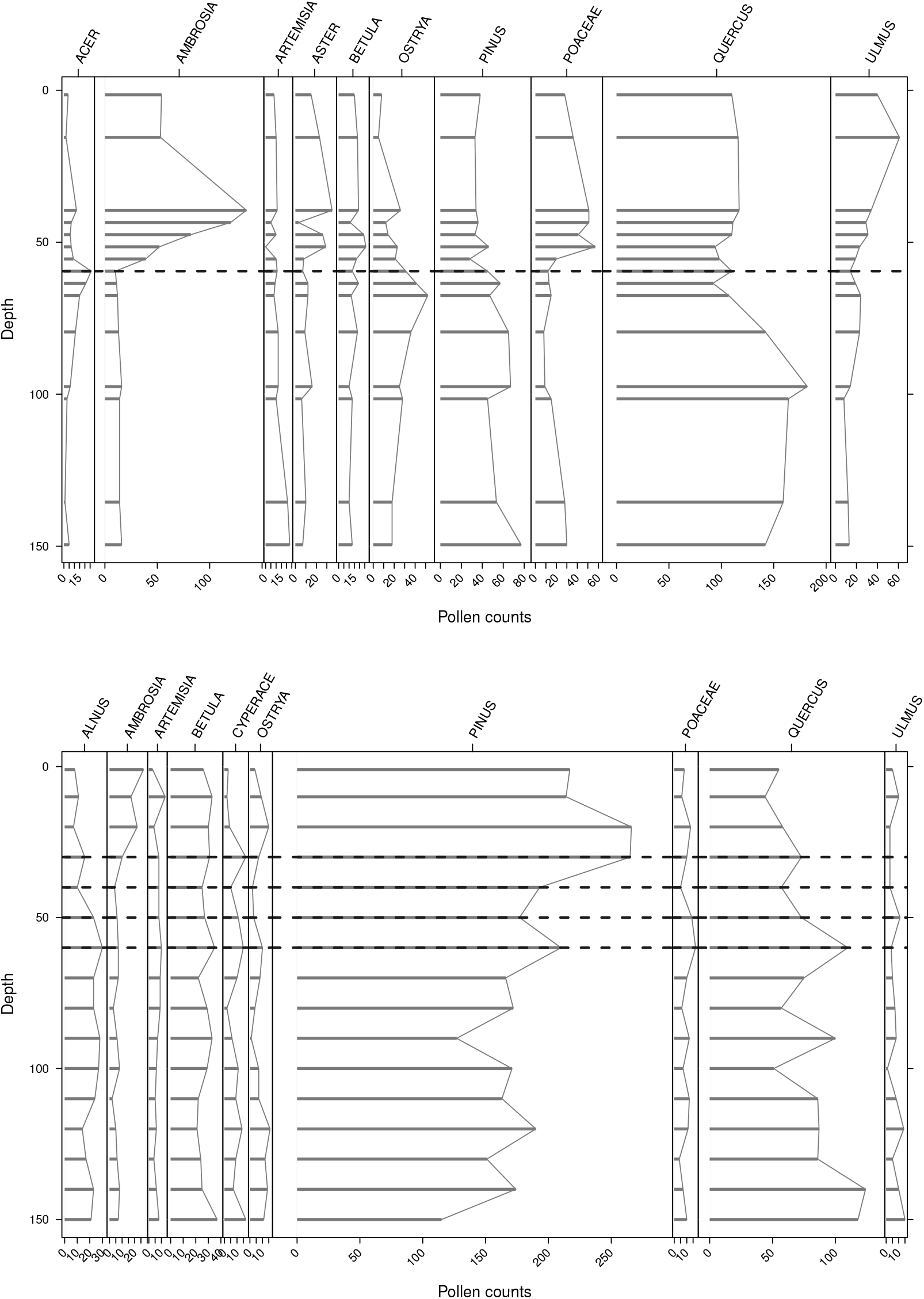
Pollen diagrams of pollen counts as a function of core depth for the ten most common taxa for Brownsby and Pennington Lake. Dashed horizontal lines indicate the expert-identified pre-settlement samples. For some sites, such as Brownsby Lake, the settlement horizon is easily identified, and all experts were in agreement with respect to its location (top panel). For other sites, such as Pennington Lake, the settlement horizon is less obvious, resulting in multiple inferred pre-settlement samples (bottom panel).

**Figure B:**
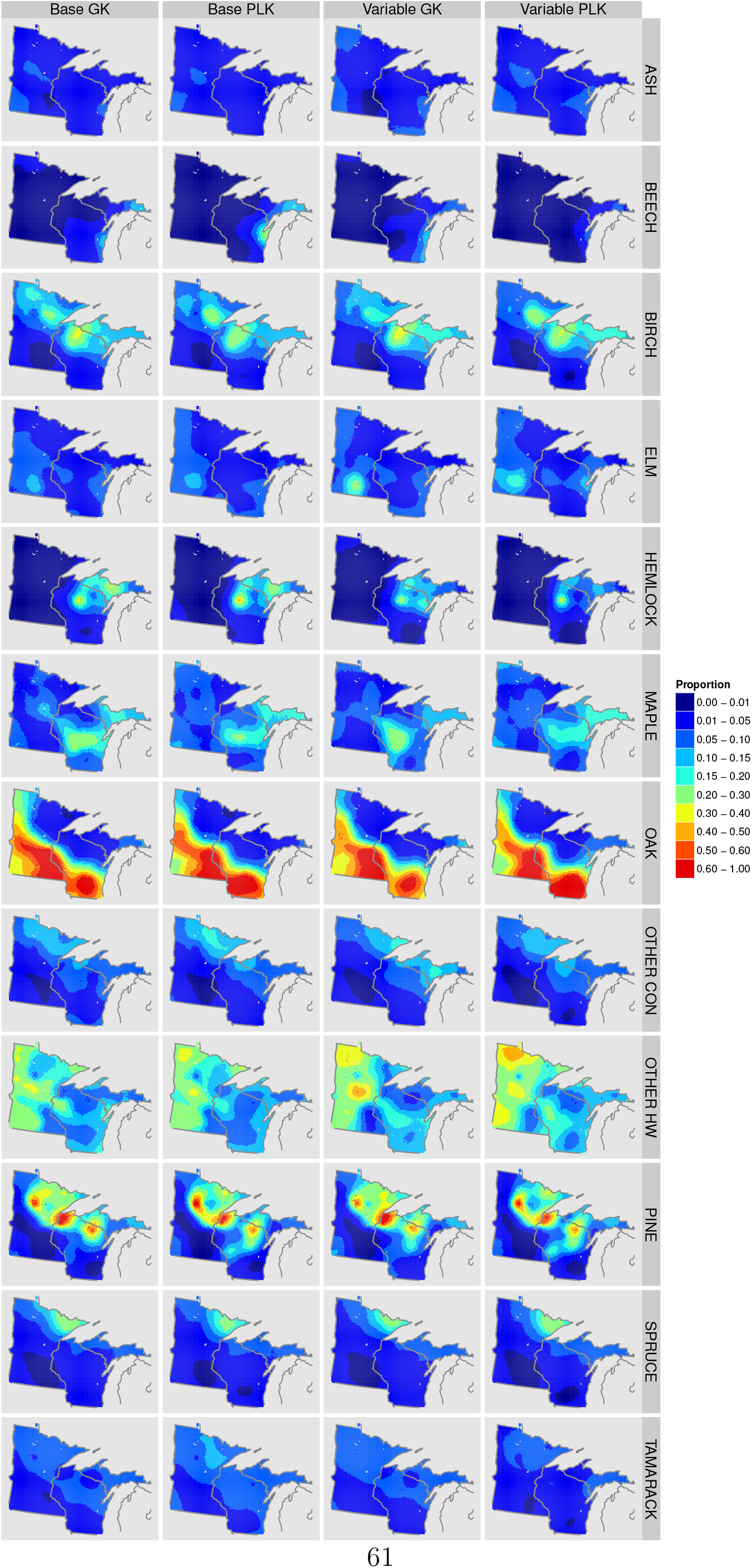
Heat maps of the STEPPS composition estimates, based on the Gaussian (GK; base and variable) and power-law (PLK; base and variable) candidate calibration models.

**Figure C:**
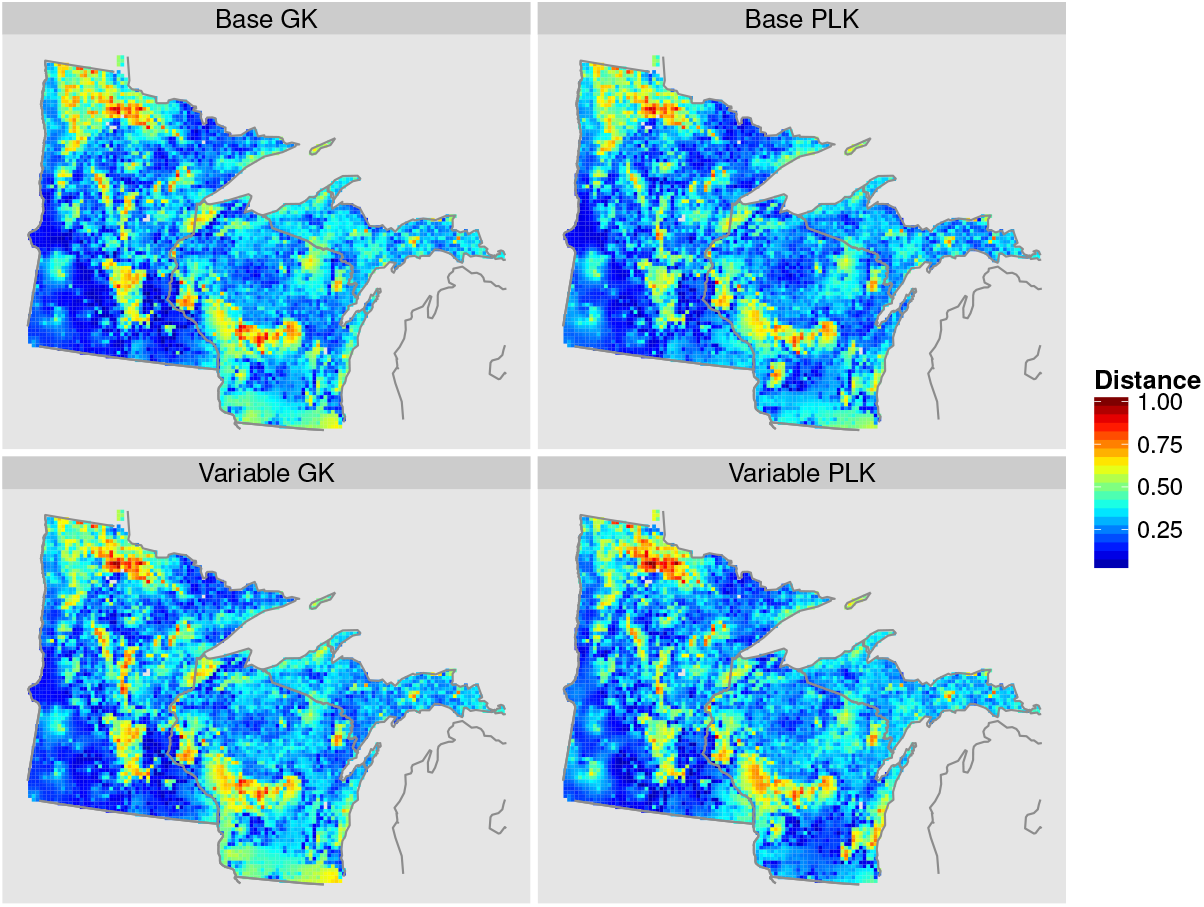
Heat maps of the dissimilarity as measured by the Euclidean distance between the PLS data and STEPPS composition estimates for different underlying calibration models.

